# Low-dose cryo-electron ptychography of proteins at sub-nanometer resolution

**DOI:** 10.1101/2024.02.12.579607

**Authors:** Berk Küçükoğlu, Inayathulla Mohammed, Ricardo C. Guerrero-Ferreira, Massimo Kube, Julika Radecke, Stephanie M. Ribet, Georgios Varnavides, Max Leo Leidl, Kelvin Lau, Sergey Nazarov, Alexander Myasnikov, Carsten Sachse, Knut Müller-Caspary, Colin Ophus, Henning Stahlberg

## Abstract

Cryo-transmission electron microscopy (cryo-EM) of frozen hydrated specimens is an efficient method for the structural analysis of purified biological molecules. However, cryo-EM and cryo-electron tomography are limited by the low signal-to-noise ratio (SNR) of recorded images, making detection of smaller particles challenging. For dose-resilient samples often studied in the physical sciences, electron ptychography – a coherent diffractive imaging technique using 4D scanning transmission electron microscopy (4D-STEM) – has recently demonstrated excellent SNR and resolution down to tens of picometers for thin specimens imaged at room temperature.

Here we applied 4D-STEM and ptychographic data analysis to frozen hydrated proteins, reaching sub-nanometer resolution 3D reconstructions. We employed low-dose cryo-EM with an aberration-corrected, convergent electron beam to collect 4D-STEM data for our reconstructions. The high frame rate of the electron detector allowed us to record large datasets of electron diffraction patterns with substantial overlaps between the interaction volumes of adjacent scan positions, from which the scattering potentials of the samples were iteratively reconstructed. The reconstructed micrographs show strong SNR enabling the reconstruction of the structure of apoferritin protein at up to 5.8 Å resolution. We also show structural analysis of the Phi92 capsid and sheath, tobacco mosaic virus, and bacteriorhodopsin at slightly lower resolutions.

## Main

Cryo-transmission electron microscopy (cryo-EM) has revolutionized life sciences and pharmaceutical research in academia and industry. Cryo-EM has been shown to be able to determine in a few hours at near-atomic resolution the structure of proteins that have been frozen as single particles in a thin aqueous layer, provided the proteins are available in sufficient concentration as homogeneous populations, sufficiently stable conformations, and embedded in random orientations in a thin ice layer without excessive adsorption to the air-water interface. Cryo-EM analysis of proteins suffers from the low signal-to-noise ratio (SNR) in the images, so that only larger protein particles typically bigger than 50 kDa, or larger details in cryo-electron tomography reconstructions of tissue slices can be analyzed at high resolution. The low SNR stems from the fact that for an electron beam, proteins are weak-phase objects and are highly fragile under the beam. Typically, the electron fluence must be limited to below 20 e^−^/Å^2^, if a protein is to be imaged at high resolution. Data can be recorded with slightly higher doses if fractionation of the electron dose is used, and recorded frames are resolution-weighted with dose-dependent frequency filters before averaging, to obtain images with higher differential contrast, *i.e.*, signal-to-noise ratio (SNR)^1^. Cryo-EM furthermore suffers from particle movements under the electron beam, which can be corrected partly during image processing through motion correction in dose-fractionated "movies"^2–4^. Finally, conventional cryo-EM suffers from the oscillating contrast transfer function (CTF), which is dampened towards higher resolution from the limited spatial and temporal coherence of the beam, the detector modulation transfer function (MTF), and non-corrected specimen movements, among others. Phase plates, such as Volta or laser phase plates^5,6^ partly improve the CTF for low-resolution components, but so far have not yet shown improvements in final resolution.

An alternative data acquisition scheme is scanning transmission electron microscopy (STEM), which uses a focused electron probe that is scanned across the sample, while electron detectors record the number of electrons scattered to a certain angle covered by the detector pixel(s) as a function of the probe position^7^. Bright-field (BF) STEM images provide only weak phase contrast^8^, and dark-field (DF) STEM yields high mass-thickness contrast, yet at low dose efficiency, so that STEM was until recently of limited use in the life sciences. STEM Z-contrast imaging, employing annular detectors covering high scattering angles, is based on Rutherford scattering, where the detector signal is approximately proportional to the square of the atomic number and linear to the number of atoms within the interaction volume. This linearity of the high-angle annular DF (HAADF) STEM signal has been exploited for detecting heavier atoms in proteins by cryo-STEM with an ADF signal^9^. It has also been exploited for mass measurements of protein particles that had been freeze-dried on ultra-thin carbon film supports^10,11^. Early attempts of high-resolution low-dose aberration-corrected HAADF STEM imaging showed only amplitude contrast and was found unsuitable for life sciences imaging^12^. However, cryo-STEM tomography, combining BF and DF STEM with sample tilt series to compute a 3D reconstruction of the vitrified specimen^13^, has shown promising results for thicker biological specimens up to 1 µm diameter. More recently, integrated differential phase contrast (iDPC) STEM was applied to cryo-EM specimens, reaching 3.5 Å resolution for protein 3D reconstructions from data recorded with a quadrant-detector^7,14^.

Beyond monolithic or segmented detectors, the use of pixelated detectors in STEM allows recording a 2D image of the diffracted probe for each 2D probe position, enabling advanced analysis in this so-called 4D-STEM method^15^. For example, electron dose-efficient recovery of the specimen’s scattering phase can be achieved by ptychographic processing of the recorded diffraction patterns^16^. For materials sciences specimens, this method has recently established resolution records in the picometer range^17,18^. Besides the inverse multi-slice schemes used in that work, which consider multiple scattering events during reconstruction but require comparably high electron doses, several other algorithms exist that employ the single-scattering projection assumption. The latter include direct methods, such as single-sideband ptychography^19,20^ or Wigner distribution deconvolution^21,22^ for 4D-STEM datasets recorded with a focused probe, or parallax reconstructions from tilt-corrected BF STEM data^23–26^ recorded with a defocused probe, to provide phase information with real-time capabilities. Alternatively, iterative methods, such as the computationally more demanding proximal gradient algorithms such as difference map calculations^27^, ePIE^28^, or ptychography reconstructions^29,30^ can be applied.

4D-STEM with an aberration-free electron beam that is focused onto the specimen typically scans the sample with a narrow Airy disk profile, limited to only a short vertical extension, depending on the convergence semi-angle (CSA) used to form the probe^7^. 4D-STEM can also be performed with a strongly defocused electron probe (**Figure 1**), so that the convergent electron beam illuminates the specimen with a disk large enough to expose an entire typical single-particle protein in the vitrified cryo-EM specimen as one single scattering experiment. This probe can then be scanned across the specimen surface at a step size in the nm range, so that diffraction patterns are recorded from probe positions that share significant overlap in the (x,y) sample plane. This overlap enables phasing of the recorded data for structure reconstruction, and it eliminates the twin-image problem or mirrored reconstructions.

**Fig. 1:**
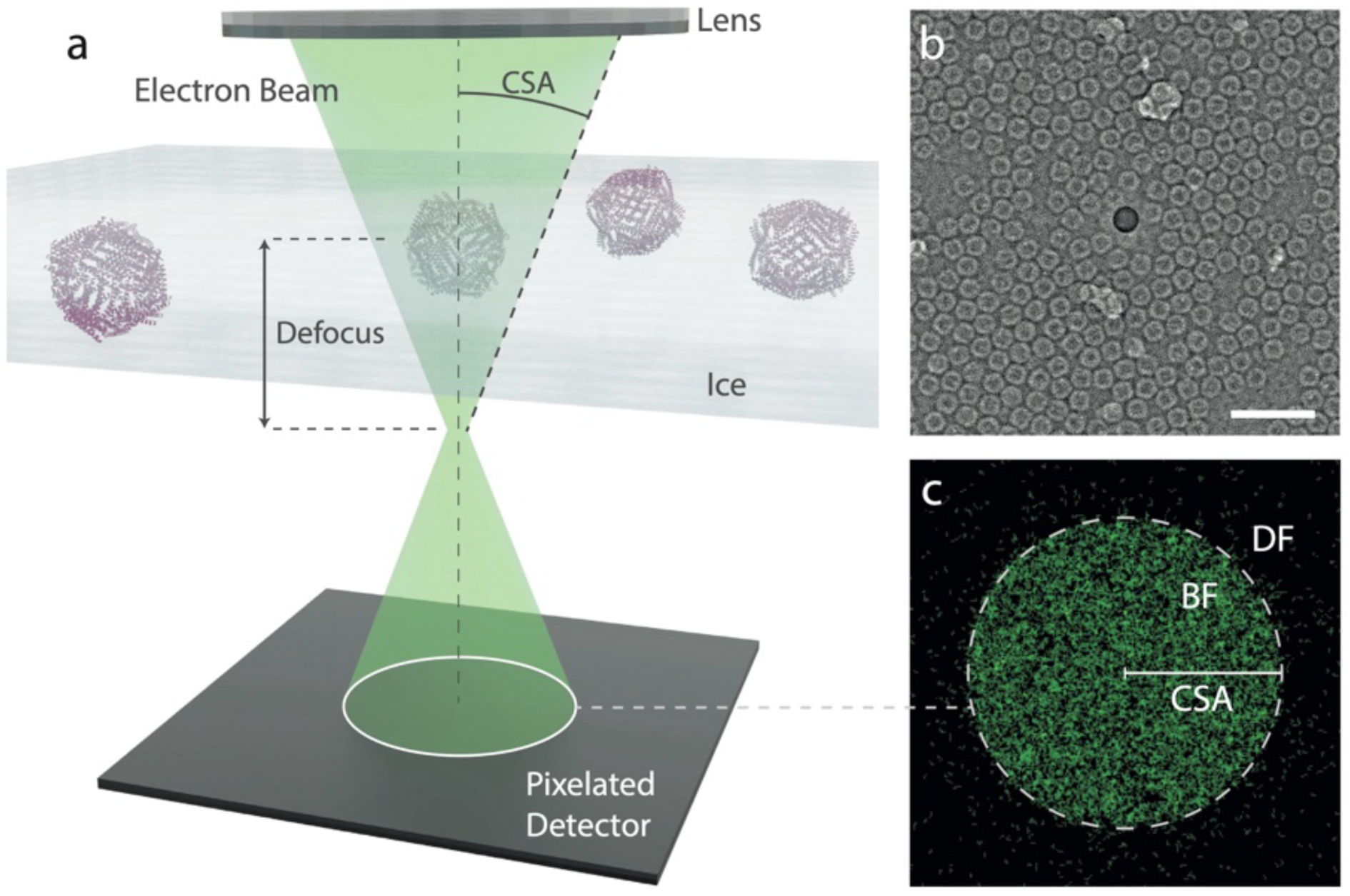
Experimental setup of 4D-STEM. **a**, A vitrified life sciences sample is illuminated with a convergent electron beam. Electrons are recorded using a pixelated detector. The convergence semi angle (CSA) is in the range of a few mrad. The beam is focused to a plane above or below sample with a defocus in the micrometer range. **b**, An example ptychographic reconstruction of apoferritin. Scale bar: 50 nm. **c**, The sum of three 4D-STEM diffraction patterns. A typical diffraction pattern shows most of the electrons in the bright field (BF) disk of the radius corresponding to the CSA of 6.1 mrad, and some electrons are scattered to higher angles to the dark field (DF) region.

4D-STEM has recently gained significant momentum from the availability of ultrafast pixelated electron detectors. Extending 4D-STEM to study beam-sensitive biological cryo-EM specimens, however, faces several experimental and computational hurdles. For low-dose experiments, scan strategies must be optimized for most efficient handling of specimen motion and specimen charging. Data processing workflows have to be established that can deal with the very low signal-to-noise ratio in the low-dose datasets and with the enormous quantity of recorded data, reaching terabytes in size for one sample. Nevertheless, significant progress in life sciences 4D-STEM has recently been demonstrated by the application of 4D-STEM ptychography to frozen-hydrated viruses ^16,31^ and virus-like particles^32^.

Here, we performed low-dose 4D-STEM studies with a defocused beam and ptychographic data processing of frozen-hydrated single particle protein samples (**Fig. 1**) and showed that structures of proteins at sub-nm resolution can be achieved. We implement cryo-electron ptychography with an aberration corrected cryo-EM instrument and fast hybrid pixel detector, and we discuss the current state of automated data collection and data processing.

### Ptychography of cryo-EM samples

Ptychographic data were recorded with an aberration-corrected, convergent electron beam as “probe” that was focused below the sample. However, as ptychography reconstructs both, the complex-valued sample exit wave and the complex-valued electron probe function including its aberrations, aberration correction is not a requirement for the work presented here, since CSAs of only 4 to 6 mrad and larger defocus values around 2 µm were used. For the 4 mrad CSA and 2 µm defocus, the probe corresponded to a beam diameter on the sample of ∼16 nm. A beam dwell time of 250 µs and a STEM step size of 2 nm, corresponding to 84% beam overlap for two adjacent circular probes, were used. Using a fluence of ∼0.5 e^-^/Å^2^ per probe position, an average total electron fluence of ∼35 e^-^/Å^2^ was applied to the sample during one scan of 128×128 positions. The low electron dose and parameters used in our experiments meant that on average less than 20% of the detector pixels received an electron for one probe position, making the recorded diffraction patterns very noisy.

A dataset of gold nanoparticles on a carbon film was collected before beginning experiments to calibrate rotation angles and CSA, and a diffraction pattern with the specimen removed from the beam path was recorded to serve as estimation for the probe-forming aperture in the iterative reconstruction algorithm. Data collection was then performed with a setup as shown in **Fig. 2**. The software SerialEM^33^ was used to control the Titan Krios instrument, take real-space TEM images of the grid, and compile a table of X,Y coordinates, indicating the specimen locations that have ice holes with suitable ice thickness and a high particle density. After this, the instrument was switched to STEM mode with a convergent beam, and a custom Python script was used for automated 4D-STEM data collection. Data collection was performed at approximately 144 specimen locations per hour, each comprising a 128×128 position 4D-STEM scan, and resulted in approximately 310 GB of raw data per hour before compression.

**Fig. 2:**
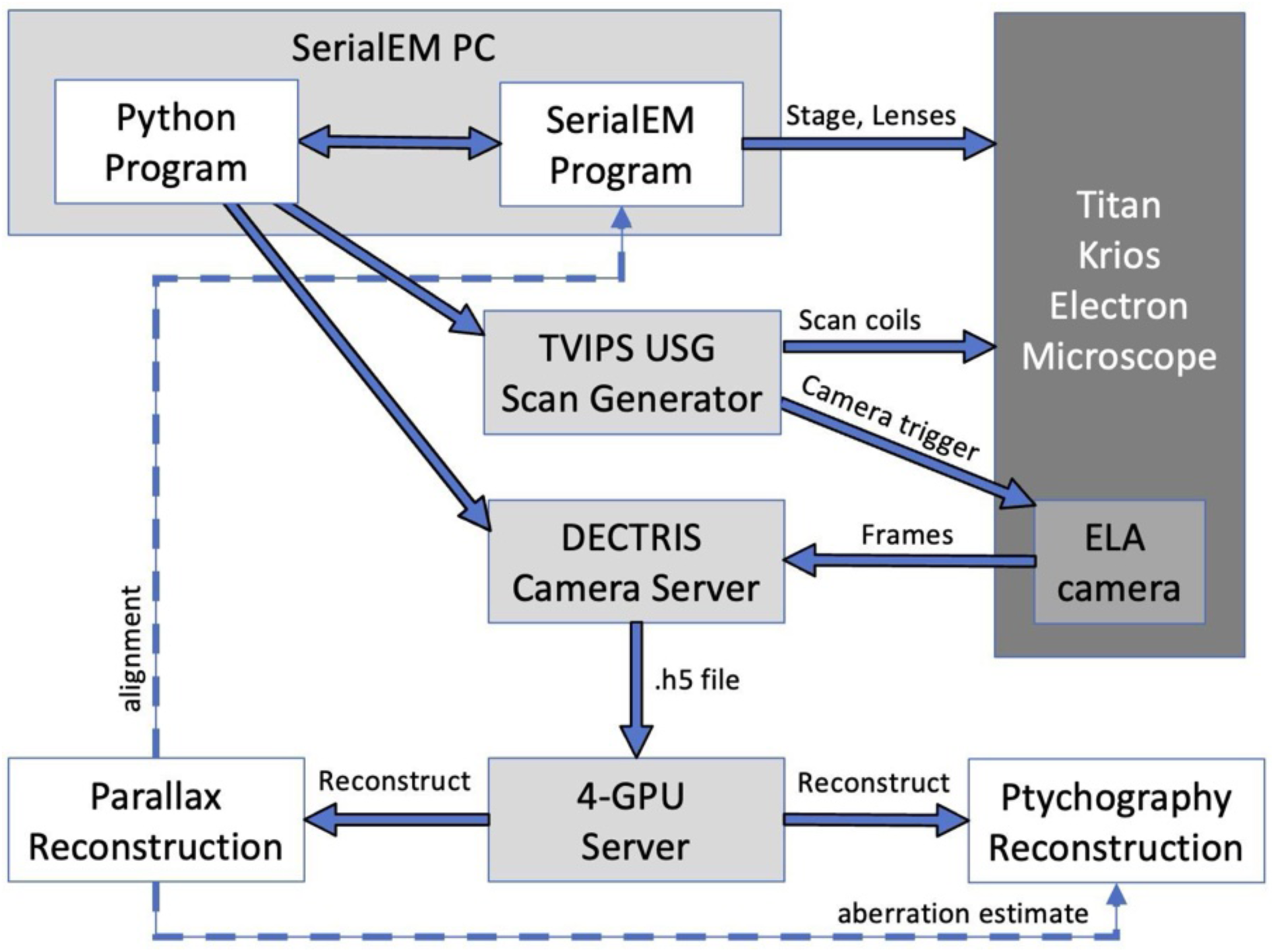
The ptychography data collection workflow. A python client running on the SerialEM Server controls the electron microscope via the SerialEM software, and coordinates the TVIPS USG scan generator, the Dectris ELA camera recording, and the multi-GPU processing of recorded diffraction patterns. Computers: grey. Software: white. Microscope: dark grey. Reconstructed images: white. Dotted line: The parallax reconstruction is used to provide on-the-fly feedback about the data collection and serve as aberration estimate.

Data collection was monitored in real-time by computing a parallax (or tilt-corrected BF-STEM) reconstruction with the py4DSTEM software^34^. This enabled fast, yet reliable, phase reconstructions to guide acquisition, and provided the probe estimate necessary for the more computationally-intensive ptychographic reconstructions. Higher-resolution ptychography reconstructions of the scattering potentials were calculated offline with py4DSTEM on a 4-GPU computer, obtaining up to 70 ptychographic reconstructions per hour.

### Structural analysis of proteins

The cryo-electron ptychography pipeline was applied to apoferritin (ApoF) protein, the phi92 bacteriophage, the tobacco mosaic virus (TMV), and 2D crystals of bacteriorhodopsin (BR). To document the outstanding quality of the ApoF protein and its suitability for high-resolution data collection, apoferritin was also imaged by conventional TEM (CTEM), before being studied using the 4D-STEM pipeline.

#### Conventional Cryo-EM imaging of Apoferritin

This spherical apoferritin particle is a protein complex of approximately 12 nm diameter with a particle weight of 485 kDa. Its octahedral 24-fold symmetry and unusually robust structural integrity make it the ideal test specimen for high-resolution assays^35,36^. Conventional TEM (CTEM), using dose-fractionated cryo-EM on a Titan Krios G4 equipped with a 300kV cold-FEG electron source and a Falcon4i direct electron detector was used to record 9,846 dose-fractionated real-space images, from which 1’899’133 ApoF particles were automatically selected and analyzed with the software cryoSPARC^37^, resulting in a 3D map from 411’705 particles at a resolution of 1.09Å (**Extended Data Figs. 2 and 3**). Data showed a ResLog B-factor of 24.5 Å^2^, indicating that the ApoF sample and the obtained cryo-EM grid preparations were of outstanding quality and not limiting resolution for a structural analysis (**Supplementary Table 1).**

The same protein sample was also used for dose-fractionated real-space cryo-EM imaging on a different Titan Krios G4, which also was equipped with a 300kV cold-FEG and a Falcon4i detector, but in addition was equipped with a Cs probe corrector above the sample. From 450 dose-fractionated real-space images, 37,592 particles were automatically selected and analyzed with cryoSPARC, resulting in a 3D map from 22,504 particles at a resolution of 1.49 Å (**Extended Data Figs. 4**). Data showed a Guinier plot B-factor of 24.1 Å^2^ (**Supplementary Table 1).**

#### Cryo-electron ptychography of Apoferritin, Phi92 bacteriophage, and TMV

4D-STEM was then used on the same ApoF grid as before with the Cs probe corrected Titan Krios, recording data with a CSA of 4 mrad. Online parallax reconstructions computed during data collection provided instant feedback on the quality of the data collection. Offline computed higher-resolution reconstructions were done by ptychography computation of the sample scattering potentials, which are obtained when assuming an amplitude function of unity and computing the resulting image only based on the reconstructed phase information^24^. Obtained 2D potential images were subjected to automated particle picking, resulting in 13’655 particles. Particle classification, alignment and 3D reconstruction using cryoSPARC^37^ resulted in a 3D map of the protein from 11’552 particles at a resolution of 5.8 Å (**Fig. 3a, Extended Data Figs. 5 and 8**, and **Supplementary Table 1**).

**Fig. 3:**
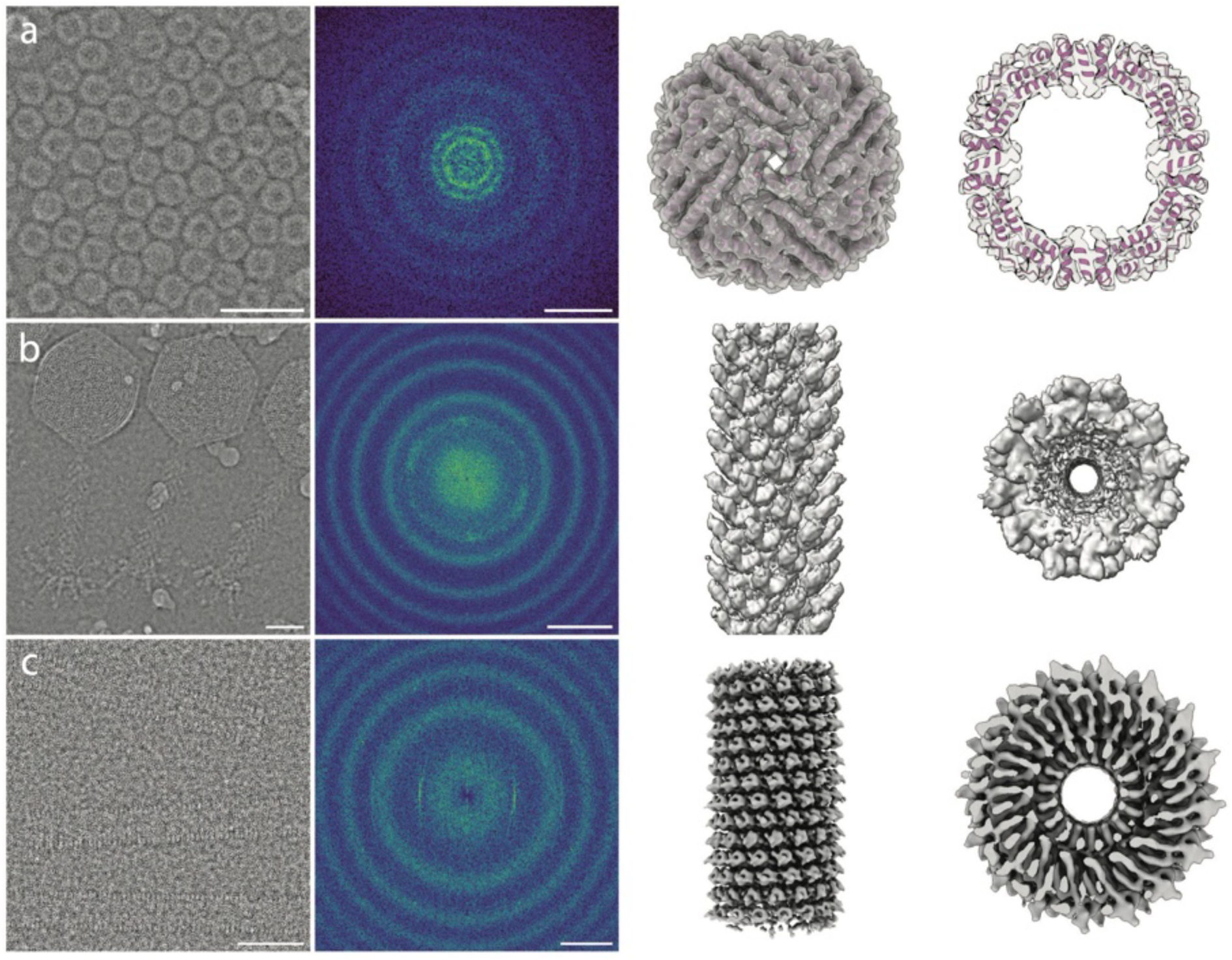
Cryo-electron ptychography reconstructions of proteins. **a**, Apoferritin. **b**, Bacteriophage Phi92 tail protein. **c**, Tobacco mosaic virus (TMV). From left to right: Ptychographic image reconstruction (Scale bars are 30 nm); The same image Fourier transformed, showing circular amplitude oscillations (Scale bars are 0.5 nm^-1^); The 3D reconstruction of the protein; A cross-section of the 3D reconstruction for **a** and a bottom view of the 3D reconstructions for **b** and **c**.

4D-STEM was also used to study the Phi92 bacteriophage, using a CSA of 5.1 mrad. Data processing, using helical reconstruction for the bacteriophage’s sheath was conducted using cryoSPARC, while the capsid reconstruction was performed using RELION^38,39^, resulting in a reconstruction of the Phi92 sheath from 1,600 particles at 8.4 Å resolution, and of the Phi92 capsid from 182 particles at 12.1 Å resolution (**Fig. 3b, Extended Data Figs. 6 and 8**, and **Supplementary Table 1**).

4D STEM was also applied to study the tobacco mosaic virus (TMV), using a CSA of 6.1 mrad. Processing of the ptychography potential maps with RELION resulted in a reconstruction from 2,120 particles at 6.4 Å resolution (**Fig. 3c, Extended Data Figs. 7 and 8**, and **Supplementary Table 1**).

#### Cryo-electron ptychography of bacteriorhodopsin 2D crystals

4D STEM was also applied to study the projection map of glucose-embedded bacteriorhodopsin (BR) 2D crystals. These single-layer phospholipid membrane protein 2D crystals have a lateral size of roughly 500 nm and a thickness of roughly 6 nm^40^. The crystals contain BR trimers that are embedded in the membrane in a P3-symmetric hexagonal lattice of 6.3 nm lattice spacing (120° included angle). A 4D STEM dataset was recorded from 256×256 positions at 4 mrad CSA with a total dose of 19 e^-^/Å^2^. Parallax and a ptychography reconstructions were computed from this one dataset, which both showed crystal diffraction spots in their Fourier transforms (**Extended Data Fig. 9**). Crystallographic processing of these two reconstructions with the FOCUS software^41^ allowed the reconstruction of the projection maps at around 6 Å resolution from each, which required for the parallax reconstruction a correction of the oscillating CTF by Wiener filtration, while the ptychography map allowed reconstruction of the correct projection map without any CTF correction. The intensity of the diffraction spots above background, and the intensity of the background intensity in the vicinity of the diffraction spots was evaluated in FOCUS, showing a faster decay (*i.e.*, stronger negative B factor) of intensities for the parallax reconstruction than for the ptychography reconstruction (**Extended Data Fig. 9**, and **Supplementary Table 1**).

## Discussion

Semi-automated 4D-STEM was implemented as a cryo-electron ptychography pipeline and applied to several biological test specimens. For reconstructing micrographs from recorded datasets, the single-slice ptychography reconstruction implemented in the py4DSTEM software was used. This algorithm refines the estimate for the *Probe* and the *Object* from the measured intensities in the recorded diffraction patterns as amplitude constraints in Fourier space, and by exploiting the knowledge about the position overlap in real space. The *Probe* and *Object* are iteratively refined, using a stochastic gradient descent algorithm. In the initiation of the projection algorithm, an empirically determined hyper-parameter tuning is also integrated, which improves convergence^24^. For details, see the **Online Methods**.

For CTEM imaging, the contrast transfer function (CTF) of the instrument is well characterized and its correction provides access to the highest resolution, including correction of higher-order aberrations^39^. Leidl et al. (2023) recently demonstrated that for thicker biological specimens, multiple electron scattering needs to be taken into account if structural data significantly beyond 1.2 Å resolution were to be recovered^42^.

For 4D-STEM, the CTF is dependent on the reconstruction algorithm as well as acquisition parameters used.

For ptychography reconstructions, computed Fourier transforms of the 2D potential maps obtained by computing ptychography phase-only reconstructions, while fixing the amplitudes to constant “1”, show for 4D datasets recorded with a defocus of 2 µm the presence of ring like amplitude oscillations (**Fig. 3**), and the width of the rings is wider (with more narrow dark gaps) when data closer to focus are recorded (**Extended Data Fig. 9**). From these ptychography potential maps, a correctly phased reconstruction can be obtained without CTF correction^16,30,43^.

In contrast, for parallax reconstructions, Thon-ring amplitude oscillations are present (**Extended Data Fig. 9**), which require CTF correction including phase flipping to give access to correct structural data (**Extended Data Fig. 10**).

Electron ptychography has established resolution records for materials sciences specimens, while the algorithms for signal reconstruction and our understanding of the mechanism of contrast formation are still under active development. For life sciences samples, cryo-electron ptychography promises to also give access to the highest resolution and contrast. The absence of an oscillating or zero-crossing CTF in ptychography is highly attractive for tomography data collection, where the reconstruction of unique features from single datasets is of interest. Significant technical challenges remain that should be addressed to further improve the resolution of the method. These include the implementation of dose fractionation during data collection, which further reduces the signal-to-noise ratio of the collected diffraction pattern; reconstructed sub-frames should then be resolution-weighted and averaged, as commonly done in real-space cryo-EM^1,2^. Further required optimizations concern the level of scan precision (noise in the scan coordinates), data collection speed (camera speed and data transfer speed), post-processing algorithms under the very low dose constraints for life sciences specimens, and the extension to tomographic 3D reconstructions in combination with multi-slice algorithms^24,44^.

The frozen hydrated preparations of the proteins apoferritin, phi92 phage, and TMV studied here by 4D-STEM and ptychography resulted in 3D reconstructions at sub-nm resolutions from small datasets. Nevertheless, this resolution is still worse than what is achievable by conventional cryo-EM from similarly sized datasets (**Extended Data Fig. 4**). Resolution improvements in life sciences 4D-STEM are expected from collecting data with a larger CSA. This, however, causes a lower tolerance for sample height variations when collecting large datasets, making data collection and data processing more challenging. The CSA of 4 to 6 mrad was chosen here as a compromise. We also applied 4D-STEM without correcting for the resolution-limiting effects of specimen beam damage or beam-induced particle motion, and we did not correct for the CTF of the ptychography method. Even if the ptychography CTF does not show contrast reversals, correction of its varying amplitude would still be required to fulfill the assumptions of equal noise contributions across all resolution ranges for the maximum likelihood processing underlying the cryoSPARC and RELION software packages. Dose fractionation and dose-dependent frequency weighting, combined with correction for particle movement, and a CTF correction that normalizes the noise profile, would all increase the resolution performance of the 4D-STEM ptychography method.

The strong SNR of cryo-electron ptychography evident in the reconstructed images, the trustworthiness of the projections obtained from the purely positive CTF, and the potential of multislice reconstructions to produce a volumetric reconstruction with modest depth resolution, suggests that 4D-STEM combined with cryo-electron ptychography tomography bears great potential for 3D reconstructions of biological tissue samples. Combined with tilting, the out-of-plane information in multi-slice ptychography results in an improved Crowther Criterion measure, scaling with the number of “super-slices” used^45^. For cellular samples, where the uniqueness of the sample does not allow the averaging of signal from multiple particles, and where thicker slices of the sample are often of interest, the high SNR of cryo-electron ptychography is especially promising to gain insight into the structural arrangement of smaller structures in cells and tissue.

## Methods

### Protein expression and purification

For apoferritin, the plasmid encoding the mouse heavy chain apoferritin was graciously donated by Masahide Kikkawa^46^. The plasmid was transformed into *E. coli* BL21 (DE3) cells. A saturated overnight culture was inoculated into 500 mL LB media containing Kanamycin, and grown at 37 °C and induced with 1 mM IPTG for 3 hours after reaching an OD600 of 0.6. The pellet was harvested at 5000 x g and resuspended in 20 mL Buffer A (300 mM NaCl, 20 mM HEPES 7.5, 10 mM EDTA), lysed by sonication and then centrifuged for 30 minutes at 20,000 x g at 4 °C. The supernatant was collected and heated at 70 °C for 10 minutes in a 50 mL tube on a heatblock until milky. The supernatant was mixed by inversion and heated for 5 more minutes. The mixture was centrifuged for 30 minutes at 20,000 x g at 4 °C. The supernatant (∼20mL) was precipitated with 6.65 g of ammonium sulfate, stirring at 4 °C for 10 minutes. This corresponds to ∼50-55% ammonium sulfate saturation at 20 °C. The mixture was centrifuged for 30 minutes at 20,000 x g at 4 °C. The pellet was then resuspended in 2 mL of Buffer A and dialyzed with a 12-14 kDa membrane in 2 L of Buffer A but containing only 2 mM EDTA. The sample was further purified by size exclusion chromatography and loaded directly onto a Superose 6 10/300 GL column with a mobile phase of 150 mM NaCl, 10 mM HEPES 7.5. Peak fractions that corresponded to 24-mer-sized apoferritin were pooled and reinjected for a second round of size exclusion chromatography. Peak fractions were concentrated to 8 mg/mL, using a 100 kDa concentrator and frozen at -80 °C.

The bacteriophage Phi92 sample, initially obtained by Petr Leiman (Texas University Medical Branch, TX, USA), was a kind gift from Davide Demurtas (EPFL, Lausanne, Switzerland).

The tobacco mosaic virus (TMV) sample, initially obtained by Ruben Diaz-Avalos (La Jolla Institute for Immunology, CA, USA), was a kind gift from Philippe Ringler (University of Basel, Switzerland).

The bacteriorhodopsin (BR) 2D crystal sample was a kind gift from Richard Henderson (MRC, Cambridge, UK).

### Real-space cryo-EM data collection and processing of a large ApoF dataset

3 µL aliquots of the apoferritin at 8 mg/ml were applied onto Ultrafoil gold 1.2/1.3 grids, and plunge frozen in liquid ethane with a Leica EM GP2 plunger (4 °C, 100% rel. humidity, 30 s waiting time, 3 s blotting time).

For conventional (real-space) cryo-EM, data were collected in automated manner using EPU v.3.0 on a cold-FEG fringe-free Thermo Fisher Scientific (TFS) Titan Krios G4 TEM, operating at 300 kV with aberration-free image shift (AFIS), and recording data with a Falcon 4i (TFS) electron counting direct detection camera in electron counting mode. The data were collected at a nominal magnification of 250 kx (pixel size is 0.3 Å at the specimen level) and a total dose of 40 (e^-^/Å^2^) for each exposure. The data from the 1.5 s exposures were stored as electron event recording (EER) files. The data from 3 × 3 neighboring ice holes in the carbon films were collected using beam and/or image shifting, while compensating for the additional coma aberration (AFIS). The data were collected with the nominal defocus range of −0.3 to −0.9 µm. With an average throughput of 922 images per hour, a total number of 9,846 movies were collected within one 12-h session.

In total, 9,846 movies were imported into cryoSPARC v4.4.1 (eer_upsamp_factor=2, eer_num_fractions=80). Upsampled EER movies were drift-corrected and dose-weighted with Patch Motion Correction (res_max_align=4, output_fcrop_factor=1/2), and the contrast transfer function (CTF) was estimated with Patch CTF Estimation (amp_contrast=0.07, res_max_align=3, df_search_min=1000, df_search_max=20000).

After rejection of bad images based on CTF Fit resolution (2 - 5 Å) and Relative Ice Thickness (1 - 1.2) parameters, 8,609 images were used for further processing. Initial particle picking was performed on a subset of 50 images with Blob Picker, using ring blobs of 110-120 Å diameters, NCC score of 0.3, Power Score of 261–712, and a minimum relative separation distance (diameter) of 0.5. Picked particles were extracted with a box size of 720 pixels, Fourier cropped to 120 pixels (binning 6×6 times) and subjected to the streaming reference-free 2D classification.

Two best 2D class averages were selected as templates for the Template Picker, using a particle diameter of 126 Å and an angular sampling of 10 degrees. In total, 1,899,133 particles were picked. Picked particles were inspected with Inspect Particle Picks, and the best 1,104,665 particles were extracted with a box size of 720 pixels, Fourier cropped to 120 pixels, and subjected to the 3 rounds of reference-free 2D classification and Heterogeneous Refinement. A small subset of the 5,005 particles was used for the ab-initio structure building with 3 classes.

The best 544,845 particles were re-extracted with a box size of 800 pixels and further refined with Homogeneous Refinement to the resolution of 1.21 Å (B-factor = 23 Å^2^). Custom parameters of Homogeneous Refinement included the application of octahedral (O) symmetry, per-particle defocus optimization using a Defocus Search Range of 4000 Å, per-group CTF parameter optimization with fitting the beam tilt, beam trefoil, spherical aberration, anisotropic magnification, and correction for the curvature of the Ewald sphere.

Refined particles were split into 9 optical groups with Exposure Group Utilities using Action “split”. Grouped particles were further refined with Homogeneous Refinement to the resolution of 1.15 Å (B-factor = 19 Å^2^). Custom parameters of Homogeneous Refinement include application of octahedral (O) symmetry, an initial low-pass filter of 15 Å, a dynamic mask threshold of 0.3, a dynamic mask near value of 4 and far value of 8, defocus refinement, global CTF refinement, and correction for the curvature of the Ewald sphere.

Refined particles were used as an input to Local Motion correction using a maximum alignment resolution of 2 Å and a B-factor of 20 Å^2^ during alignment. Overlapping particles were removed with a minimum separation distance of 120 Å. The remaining 411,705 particles were subjected to Reference Based Motion Correction with default settings and further refined with Homogeneous Refinement (defocus refinement, global CTF refinement, and correction for the curvature of the Ewald sphere) to the resolution of 1.09 Å (B-factor = 18 Å^2^). A ResLog B-factor analysis was performed in CryoSPARC, which reported a ResLog B-factor of 24.3 Å^2^ (**Suppl. Table 1**).

An atomic model for apoferritin was re-built manually from the previously deposited model (PDB ID 8J5A) with COOT v0.9.6^47^, and iterated with rounds of refinement with Servalcat^48^ and Molprobity v4.5.2^49^. After each round of refinement, side-chains were manually inspected and adjusted in COOT with Molprobity.

### Real-space cryo-EM data collection and processing on a probe corrected instrument

To compare with the 4D STEM Data, ApoF were collected with the same instrument as used for 4D STEM work. This is a Thermo-Fisher Scientific (TFS) Titan Krios G4, operated at 300 kV, and equipped with a Cs probe corrector (Probe Cs = 0 mm). Real-space TEM images were recorded with a Falcon 4i (TFS) direct detection camera in electron counting mode at a nominal magnification of 250 kx (pixel size is 0.3153 Å at the specimen level) with a defocus range of −0.3 to −0.8 µm. Each exposure had a total dose of 28 (e^-^/Å^2^) and was stored as an electron event representation (EER) file.

A total of 1,319 EER files were acquired and imported into cryoSPARC v4.4.1 for processing. Without any up-sampling, raw EER recordings were dose-fractionated into 32 frames (with 0.875 e^-^/Å^2^ per frame) and drift-corrected. After preprocessing and contrast transfer function (CTF) estimation, a total of 913 micrographs were selected based on CTF-cutoff value 3.4 Å and ice filtering. A further subset of 450 micrographs were randomly selected to pick roughly 37,592 particles that are comparable to the low-dose ptycho-dataset of ApoF (38,425 particles). After initial 2D classification, 40 % of the particles were excluded and the remaining 22,504 particles were re-extracted and re-centered to undergo ab-initio and homogeneous refinement in cryoSPARC. Further correction of defocus, global CTF refinement and Ewald sphere curvature correction yielded a map with the FSC 0.143 at 1.70 Å. To increase the quality and resolution of the reconstruction, per-particle local drift correction was performed based on the reference map. Homogeneous refinement of these reference-based motion corrected particles improved the resolution of ApoF to 1.49 Å (B-factor = 24.1 Å2).

### 4D-STEM data collection

Ptychographic data were recorded with the same probe-corrected TFS Titan Krios G4 as above, equipped with a cold FEG electron source operated at 300kV, TVIPS Universal Scan Generator (USG), Falcon 4i camera (TFS), and a Dectris ELA hybrid pixel detector. The microscope was operated in STEM mode with a CSA of 4 to 6 mrad and a defocus of ∼2 µm, resulting for the 4 mrad CSA in a beam diameter on the sample of ∼16 nm. Data were collected with a beam dwell time of 250 µs, and a STEM step size of 2 nm, corresponding to 84% beam overlap for two adjacent circular probes. Using a fluence of ∼0.5 e^-^/Å^2^ per probe position, an average total electron fluence of ∼35 e^-^/Å^2^ was applied to the sample during one scan. To avoid the physical gaps between the segments of the ELA detector, data were recorded on only one of the detector segments of 256×256 pixels. The STEM camera length was set so that the edge of that 256×256 px square was at 6.58 mrad. The low dose regime and parameters used in our experiments resulted in less than 20% of the detector pixels, on average, recording an electron at all. Dose fractionation, not employed here, could have been implemented by repeatedly scanning the same specimen location several times with an even lower dose, which would have further reduced the dose per pattern.

To improve convergence of the ptychographic reconstruction, especially under low-dose conditions, we optimized the alignment of the STEM probe before data acquisition, using the *Probe Corrector S-CORR* software by TFS on a standard gold cross-grating sample. Optimization of the aberrations up to twice the CSA that was later used for data collection, here to a semi-angle of 8 mrad, took approximately 10 minutes. We then collected a calibration dataset before beginning experiments to aid the optimization in the py4DSTEM software. Finally, a vacuum image of the probe was recorded in diffraction space to serve as an estimate for the probe-forming aperture in the iterative reconstruction algorithm.

For data collection, the software SerialEM^33^ was used to control the Titan Krios instrument and record a grid atlas at low magnification in image mode. Suitable grid squares on that atlas were manually selected, and for each of these, SerialEM was used still in conventional TEM mode with parallel illumination, to record grid square maps. These maps were then used to semi-automatically compile a table of X,Y coordinates, indicating the specimen locations where ice holes had suitable ice thickness and a high particle density. After this, the instrument was switched to STEM mode with a convergent beam, and a home-built Python script was used for automated 4D-STEM data collection, performing the following steps:

1. Controlling the microscope via SerialEM commands to position the instrument specimen stage.
2. Preparing the ELA detector for capturing frames.
3. Activating the TVIPS scan generator for acquisition.
4. Compressing and saving the recorded diffraction patterns for further analysis.

Data collection was performed at approx. 144 specimen locations per hour and resulted in approx. 310 GB of raw data per hour before compression. Data collection could be further accelerated by recording larger ptychography scans at each stage position, and by further accelerating the camera recording and data storage speeds, which all would have increased the amount of data recorded.

### Data analysis

Data collection was monitored in real-time by computing a parallax reconstruction^24^ with the py4DSTEM software^34^. In addition, a higher resolution ptychography reconstruction was calculated offline with py4DSTEM, with the following steps:

1. **Data loading and flat-field correction:** Electron diffraction patterns from one scan in h5 format were loaded, flat-field normalized, and hot/dead pixel corrected by replacing values of identified non-functional pixels with median values computed from their neighbors. Scan metadata such as scan step size were stored separately.
2. **Organizing 4D-STEM data:** The dataset was then reorganized into a four-dimensional array based on the scanning coordinates. Binning of diffraction patterns was not performed.
3. **Extraction of key parameters:** The py4DSTEM package was then used to compute the average diffraction pattern, identify the brightfield disk, and calculate the reciprocal space pixel size.
4. **Parallax reconstruction:** A parallax reconstruction was computed to give a first reconstruction image and to estimate probe defocus. A CTF phase-flipped reconstruction was computed to serve as real-time reference image.
5. **Running ptychographic reconstruction:** Starting from the parallax reconstruction, a ptychographic reconstruction was initiated and iterated until convergence, typically reached in 30 iterations, as monitored by an L2 norm of the difference between modeled and measured diffraction intensities as error metric for single-slice ptychography. The probe function in the ptychography reconstruction was initialized based on provided aberrations (C1 in our case) for the phase, and the known amplitudes from the vacuum measurements of the probe, which were recorded for each data collection session. The probe’s Fourier amplitude was replaced after each iteration with the amplitude from the vacuum probe measurements. The phase of the probe was also regularized after each update step by fitting a low-order surface expansion (up to radial and angular order 3) and removing residuals^24^.
6. The **potential map**, obtained from a complex reconstruction with the amplitude fixed to “1”, was chosen for further protein structural analysis. See **Extended Data Figure 10** to compare different ptychography data models.
7. **Image storage:** Finally, the reconstructed micrograph was cropped if necessary and saved for subsequent analysis.

The workload was distributed across four GPUs, each processing a single dataset. This parallel approach allowed the processing of 70 datasets per hour with a 4-GPU computer.

### 4D-STEM ptychography structure determination of proteins

<u>For apoferritin</u>, ninety cryo-electron ptychography datasets (i.e., 4D-STEM scans) were acquired using SerialEM. A grid atlas was initially acquired using the Falcon4i camera, from which suitable grid squares were manually identified to compile grid square maps and identify suitable acquisition locations. The ice thickness at these locations was estimated to be thin, as they only showed single layers of clearly contrasted ApoF particles. At these locations, automated ptychography scans with a CSA of 4 mrad were started, using an average total electron fluence of ∼35 e^-^/Å^2^ per scan.

Ptychography reconstructions of the sample potential maps, which are obtained when assuming an amplitude function of unity and computing the resulting image only based on the reconstructed phase information, were obtained at a pixel size of 1.5 Å/px. These potential maps were imported as MRC image files into cryoSPARC^37^. Micrographs were subjected to automated particle picking, resulting in 38,425 particles (**Extended Data Fig. 4**). 2D classification identified 10 classes. The class averages were used as templates to refine the automated particle picking, resulting in 13,665 particles. After another 2D classification and selection of the best classes, a total of 11,552 particles were used for ab-initio 3D structure determination. No further correction for any remaining contrast transfer function was applied.

The resulting 3D map was analyzed by gsFSC with the 0.143 cutoff criterion, showing a final resolution of 5.8 Å of the map (**Fig. 3a**, and **Extended Data Figs. 5 and 8**, and **Supplementary Table 1**).

<u>For the Phi92 bacteriophage</u>, eighty-three 4D-STEM datasets were acquired using the same methodology as before, but at a CSA of 5.1 mrad. The micrographs were recorded at a total electron dose of approximately 49 e^-^/Å². The pixel size of the final micrographs was 1.44 Å/px.

The 2D ptychographic micrographs obtained were imported as MRC image files into both cryoSPARC^37^ and RELION^38,39^ for subsequent analysis. The helical reconstruction process of the bacteriophage’s sheath was conducted using cryoSPARC, while the capsid reconstruction was performed using RELION.

Initial manual particle picking of the bacteriophage sheath in cryoSPARC yielded 2,420 particles. Following 2D classification and selection of the best classes, 1,600 particles were utilized for ab-initio 3D reconstruction. The helical parameters employed in this process included a helical rise of 35.30 Å and a twist of 26.3 degrees. The final processing stage involved non-uniform refinement (NU-Refine) with a Z fraction set at 50%, resulting in an 8.4 Å resolution map of the Phi92 sheath.

Concurrently, manual particle picking of the capsid in RELION resulted in the identification of 356 particles. After a similar 2D classification process, 182 particles were selected for the final 3D reconstruction. This approach yielded a 12.1 Å resolution map of the Phi92 capsid (**Fig. 3b**, **Extended Data Figs. 6 and 8**, and **Supplementary Table 1**).

<u>For the tobacco mosaic virus (TMV)</u>, forty-three 4D-STEM datasets were acquired, following the same protocol as before, but with a CSA of 6.1 mrad. The datasets were acquired at a total dose of approximately 32 e^-^/Å² per micrograph, and with a pixel size of 1.15 Å/px.

These 2D ptychographic micrographs were imported as MRC image files into the RELION software. The initial phase of manual particle picking of TMV in RELION resulted in 3,400 particles. Subsequent 2D classification and the selective process for the best classes reduced the number to 2,120 particles, which were then used for the ab-initio 3D reconstruction. Using helical parameters for the processing of a helical rise of 1.38 Å and a twist of 22.036 degrees resulted in a 6.4 Å resolution map of the TMV (**Fig. 3c, Extended Data Fig. 7 and 8**, and **Supplementary Table 1**).

<u>For the bacteriorhodopsin (BR) 2D crystals</u>, one single 4D-STEM dataset was acquired, following the same protocol as before, but with a CSA of 4.0 mrad. The dataset was acquired at a total dose of approximately 19 e^-^/Å² per micrograph, and with a pixel size of 0.98 Å/px.

The 2D parallax reconstruction (without prior CTF correction) and the 2D ptychographic reconstruction were imported as MRC image files into the FOCUS software^41^. The entire processing and evaluation was performed automatically in FOCUS. For the parallax reconstruction, the programs gCTF or CTFFIND3 were used to determine the defocus of 1.098 µm. For the ptychography reconstruction, no CTF was determined. The crystal lattice and crystal spots were automatically detected. The crystal lattice was computationally straightened by warping (unbending) the real-space image to yield sharper diffraction spots. Amplitudes above background, as well as phases for each spot, and the background amplitudes around each spot were measured. These were used for generation of a projection map. The amplitudes and background values were plotted against the resolution of the spots in Excel (**Extended Data Fig. 9**, and **Supplementary Table 1**).

### ResLog B-factor estimation

The information content of the recorded data from conventional TEM and 4D-STEM ptychography imaging was evaluated by calculation of the FSC_0.143_ and the Guinier-plot amplitude falloff B-factor. In addition, the Reconstruction "ResLog" B-factor^35,50^ was calculated for the real-space datasets. With the latter, the number “#” of particles required to reach a resolution of "d" Angstrom, can be calculated as:

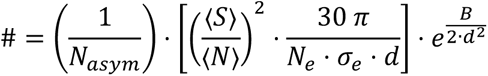

with *N_asym_* being the number of asymmetric units in the particles, 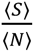 being the signal-to-noise ratio of the electrons in the recorded data, σ_*e*_ = 0.004 Å^2^ being the elastic cross-section of carbon, *N_e_* being the used electron dose, *d* being the real-space resolution in Å, and *B* being the ResLog B-factor in Å^2^.

This can be simplified to:

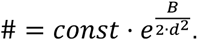

The ResLog B-factor in Å^2^ can be calculated, based on the relation:

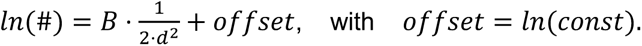

When computing the resolution *d* of a reconstruction with smaller subsets of # number of particles and plotting *ln*(#) as a function of 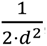 the ResLog B-factor becomes apparent as the slope of a line fitted through the experimental data points (**Extended Data Fig. 8**).

The computed FSC_0.143_, Guinier-plot B-factors, and ResLog B-factor are given in **Supplementary Table 1**.

## Data availability

For the apoferritin analysis by real-space cryo-transmission electron microscopy, the raw cryo-EM images were deposited to the Electron Microscopy Public Image Archive (EMPIAR) with accession code EMPIAR-11866. The cryo-EM map was deposited to the Electron Microscopy Data Bank (EMDB) with accession code EMDB-19436. The atomic model was deposited to the Protein Data Bank (PDB) with accession code PDB-8RQB.

For the apoferritin analysis by cryo-electron ptychography, raw data are available EMPIAR XXX and the reconstructed volume at EMD-19425.

For the Phi92 analysis by cryo-electron ptychography, raw data are available at EMPIAR XXX. The reconstructed sheath volume is at EMDB-19430 and the reconstructed capsid volume at EMD-19432.

For the TMV analysis by cryo-electron ptychography, raw data are available EMPIAR XXX and the reconstructed volume at EMD-19885.

## Code availability

The py4DSTEM package is available at https://github.com/py4dstem/py4DSTEM. The data processing Jupyter Notebook utilized here is available at https://github.com/LBEM-CH/4D-BioSTEM.

## Acknowledgements

We thank Richard Henderson for providing the BR 2D crystals, and we acknowledge fruitful discussions with Duncan Alexander, Cécile Hébert, and Marco Cantoni, EPFL Lausanne.

## Author Contributions

B.K. designed the data acquisition and processing workflow, and integrated the components of the experimental setup. B.K., I.M., and R.C.G. collected ptychography data. I.M. and B.K. performed the single particle analysis of the ptychographically acquired data. S.R., G.V., and C.O., are the developers of the py4DSTEM software and provided essential support in the processing of ptychographic data. K.L. purified and prepared the Apoferritin sample. S.N., I.M., and A.M. collected TEM images and ran Single Particle Analysis (SPA) on the Apoferritin sample. M.L.L., C.S. and K. M.-C. provided expertise and support on diffractive imaging. M.L.L and K.M.-C. Performed the analysis of power spectra of ptychographic reconstructions in Ext. Data Fig. 10. H.S. designed the experiments, provided resources, performed 2D crystal analysis, and oversaw data analysis and interpretation. B.K. and H.S. wrote the manuscript with input from all authors.

## Extended Data

**Extended Data Fig. 1:**
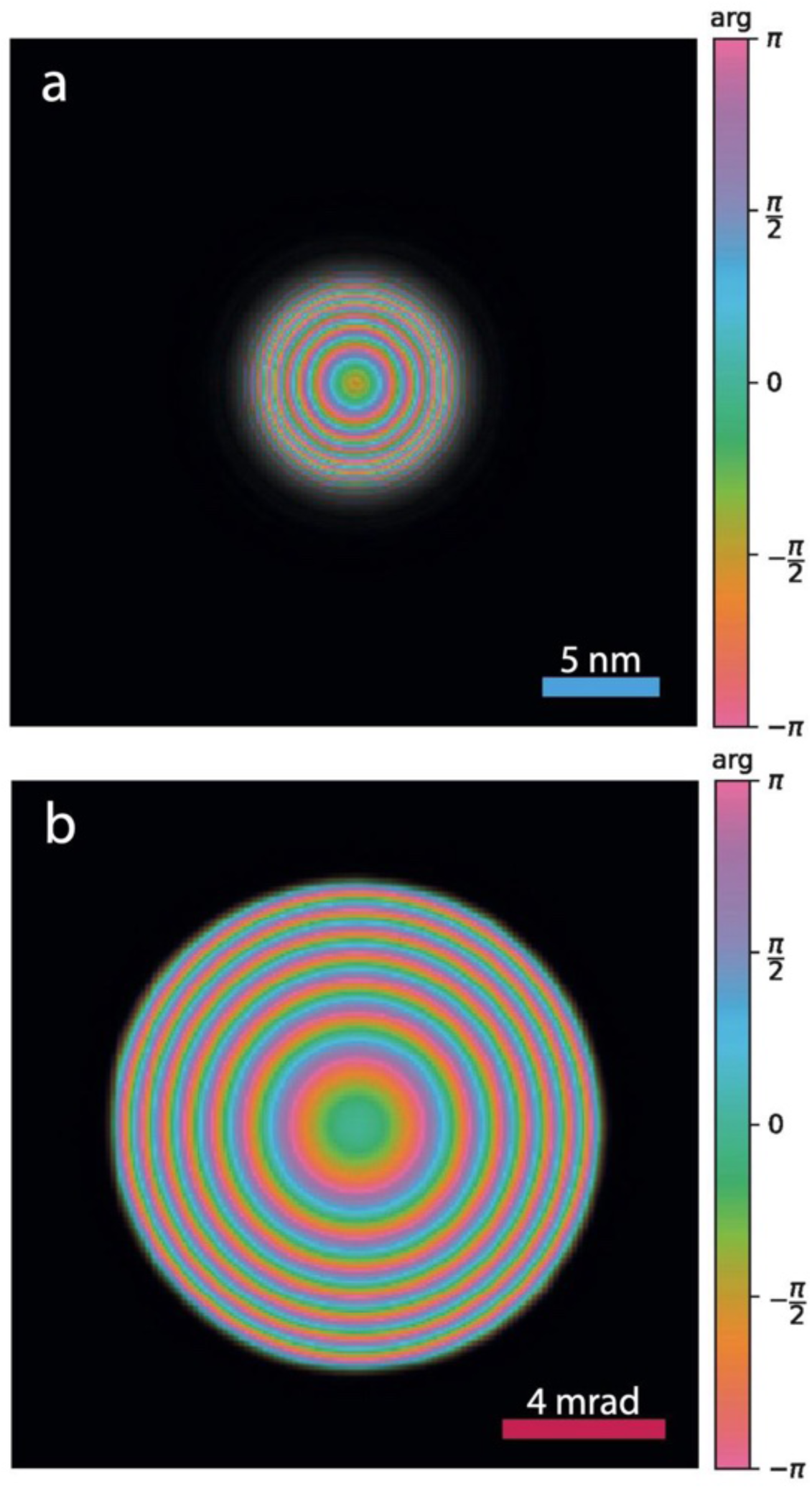
The 4D-STEM electron probe from a micrograph recorded at 6.1 mrad CSA and -758 nm defocus. a, Amplitude and phase of the reconstructed real-space electron probe on the sample. b, Amplitude and phase of the reconstructed Fourier probe on the detector.

**Extended Data Fig. 2:**
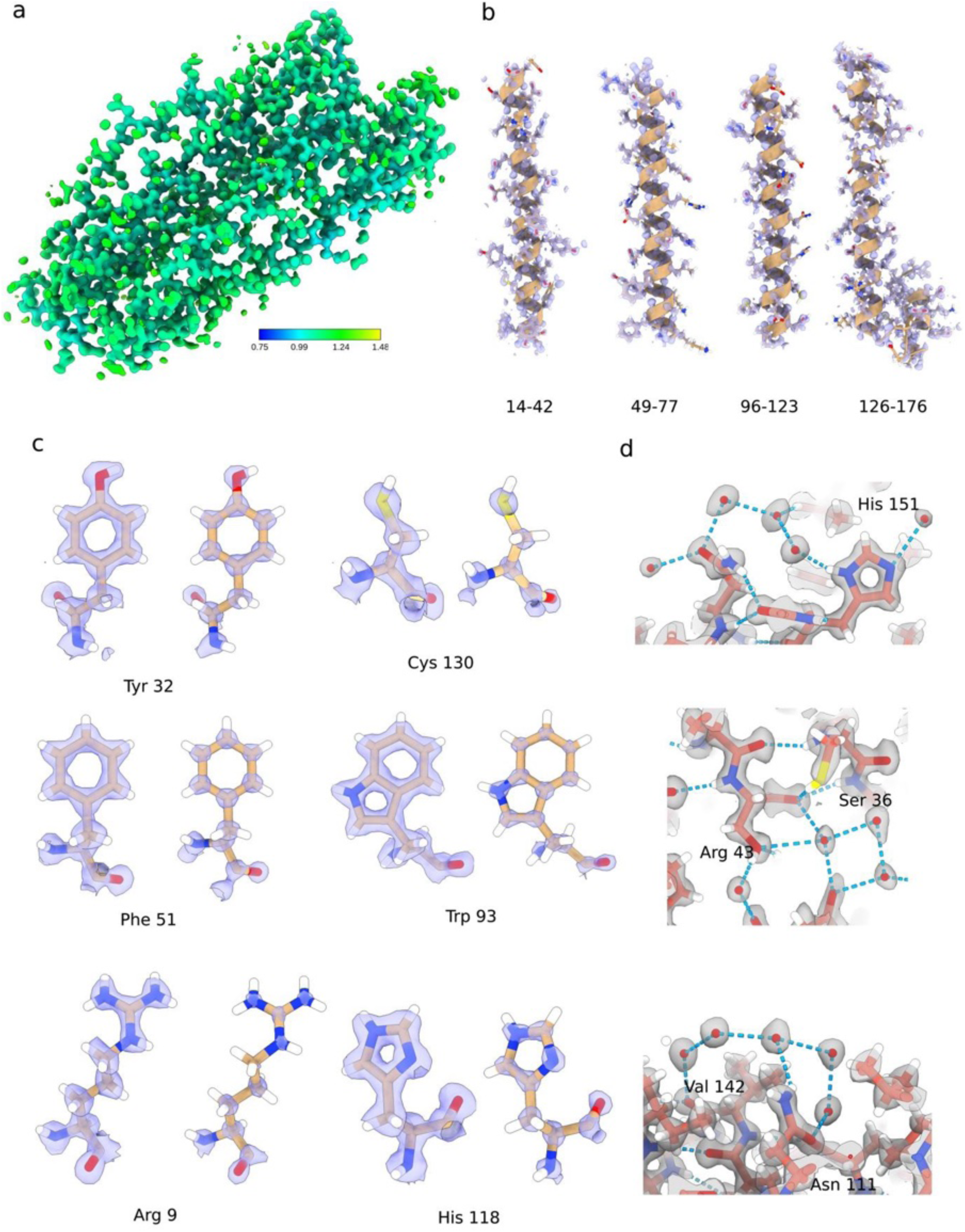
Real-space cryo-transmission electron microscopy structural analysis of Apoferritin at 1.09 Å resolution.

**Extended Data Fig. 3:**
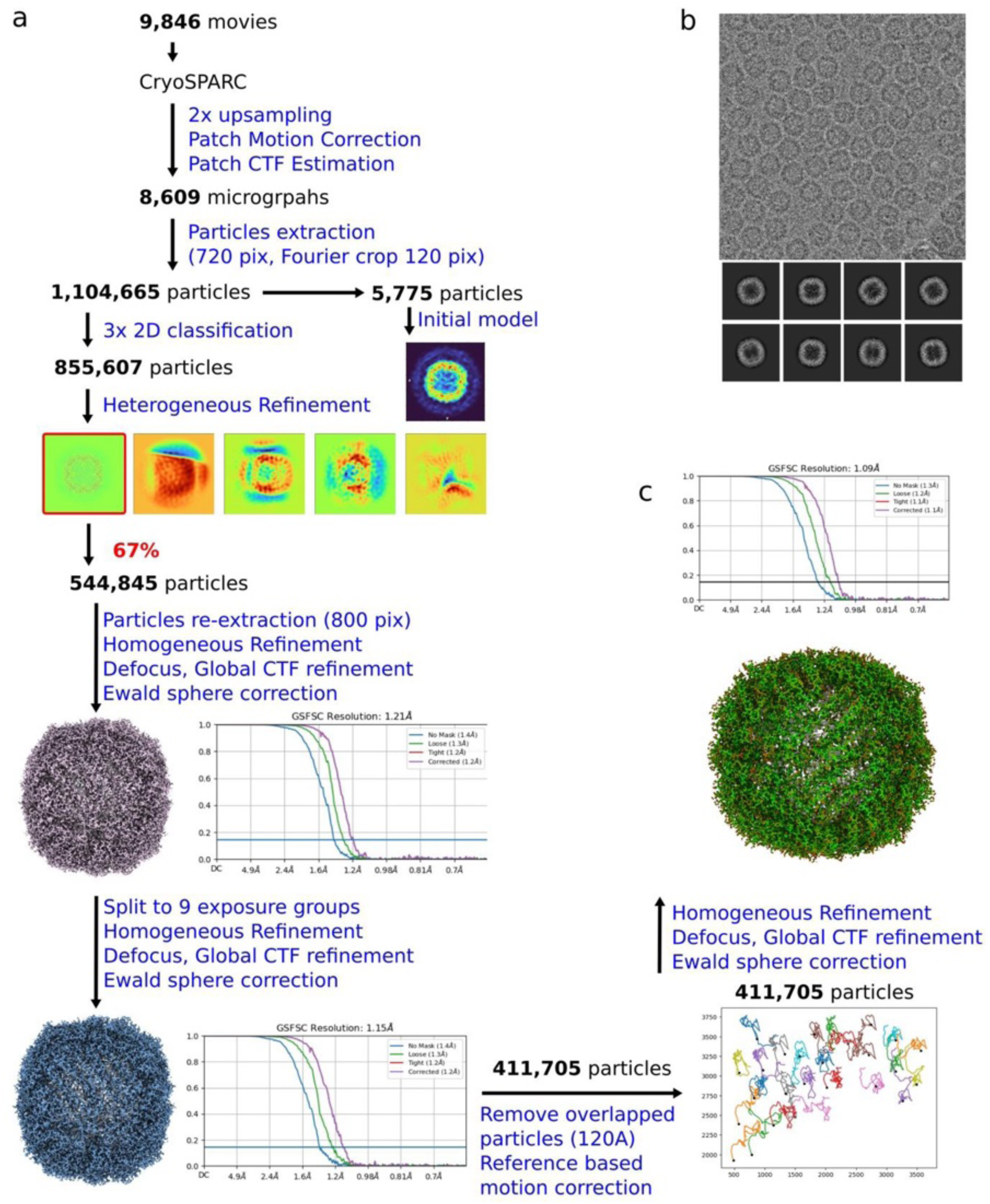
Real-space cryo-transmission electron microscopy of apoferritin and its data processing details.

**Extended Data Fig. 4:**
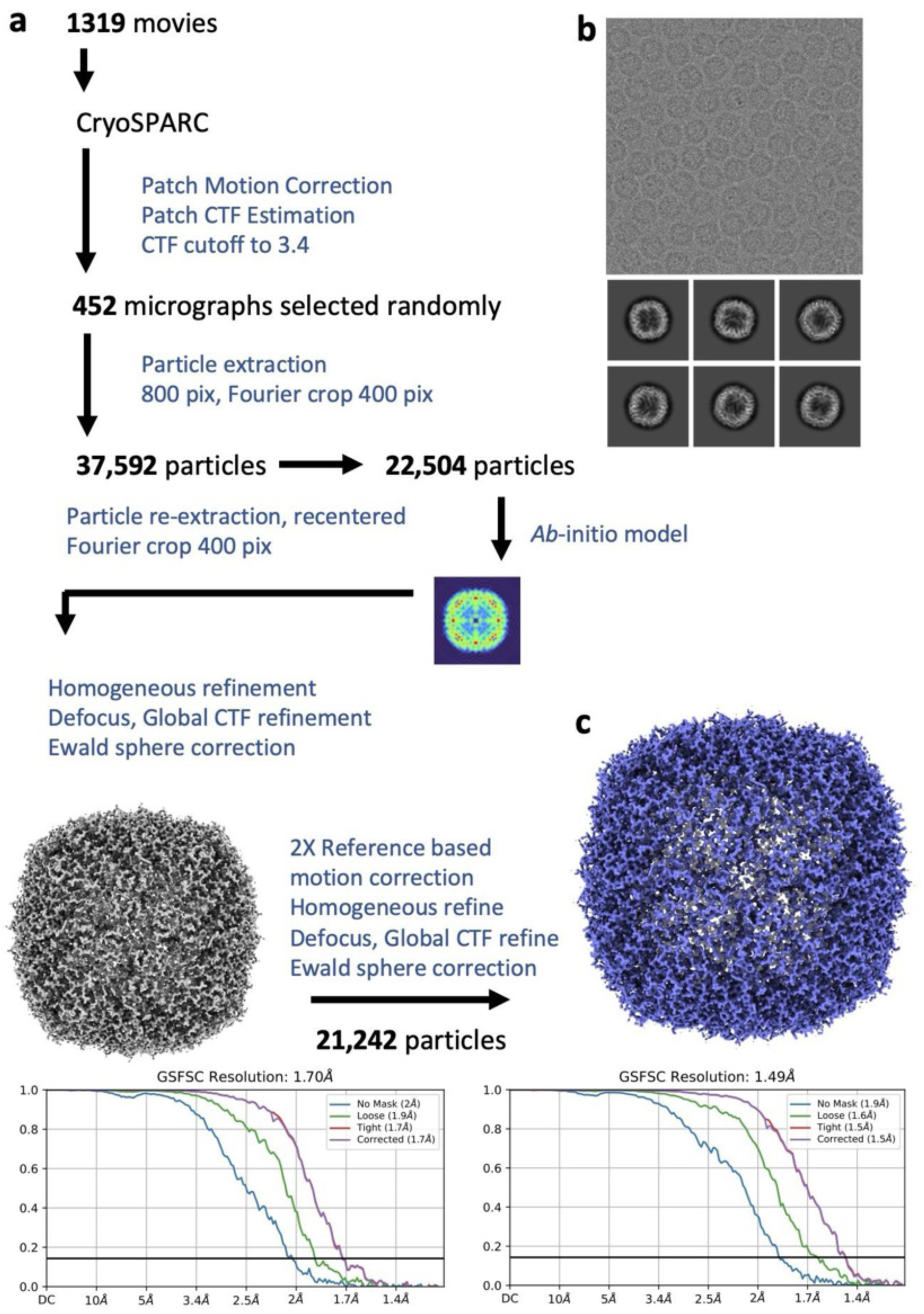
Real-space cryo-transmission electron microscopy of apoferritin, using a Cs probe corrected Titan Krios.

**Extended Data Fig. 5:**
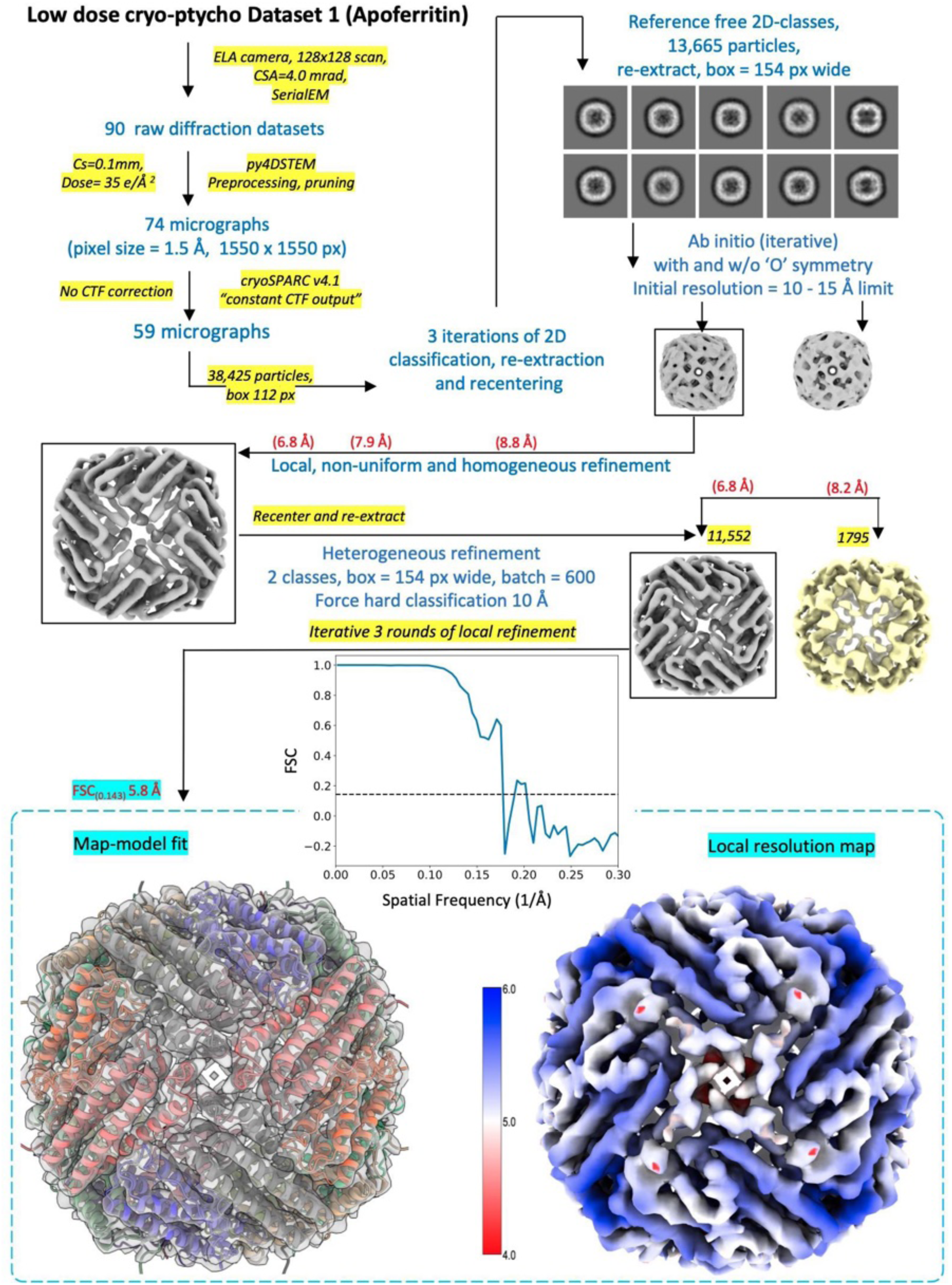
Ptychography structure determination of Apoferritin, using 4D-STEM data. The atomic model of Apoferritin used for the fitting was PDB 8J5A.

**Extended Data Fig. 6:**
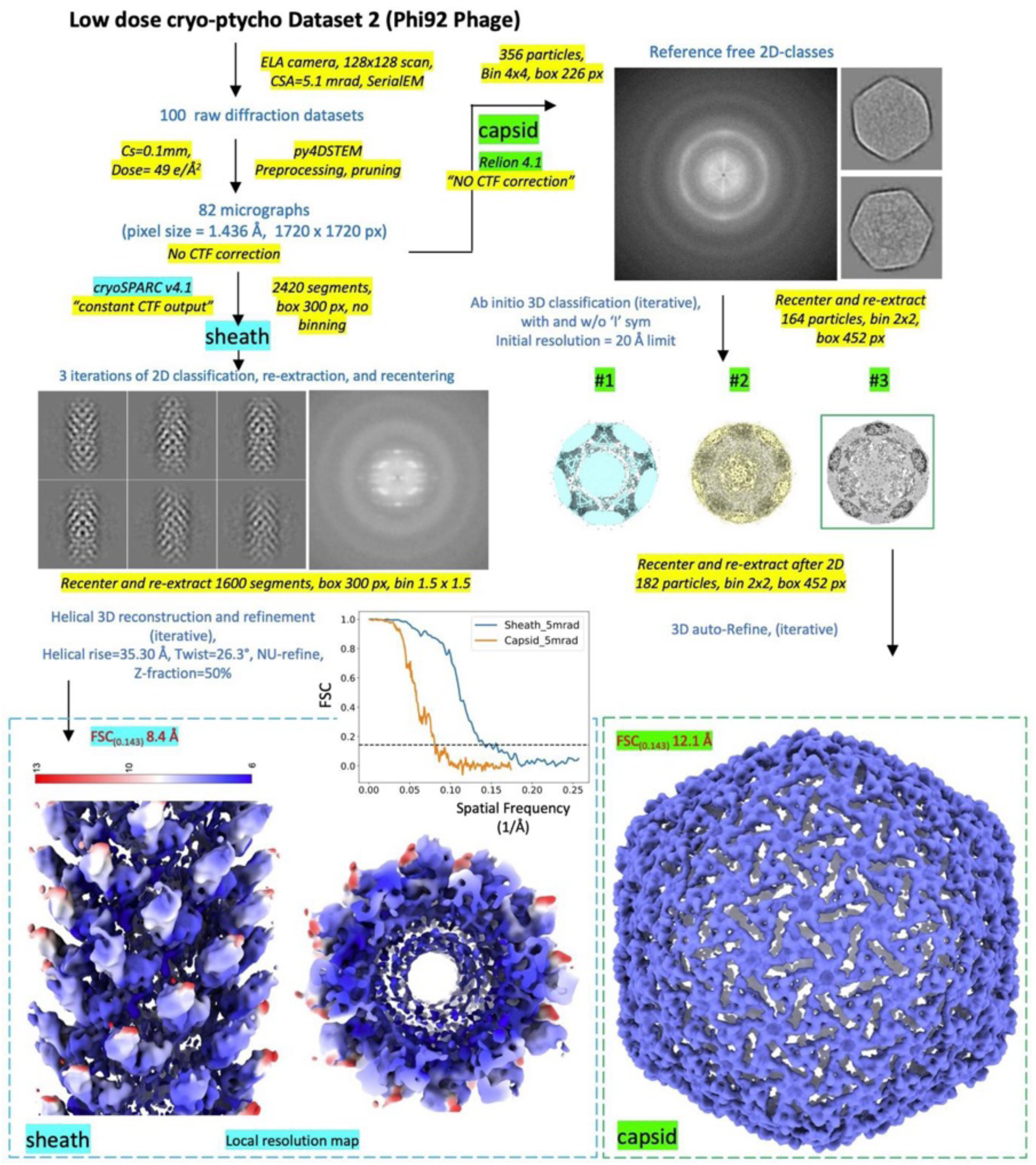
Ptychography structure determination of the bacteriophage Phi92, using 4D-STEM data.

**Extended Data Fig. 7:**
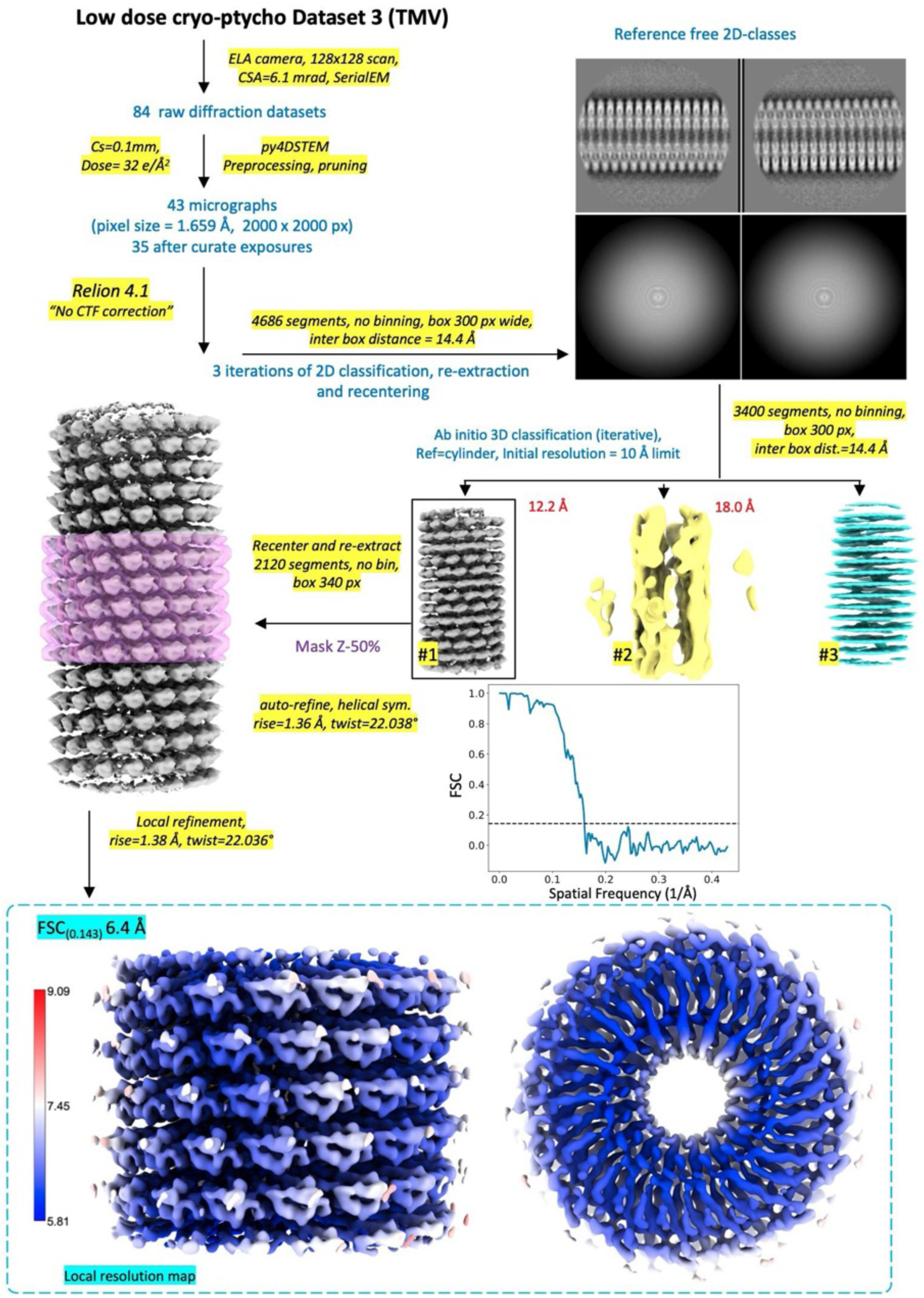
Ptychography structure determination of tobacco mosaic virus (TMV), using 4D-STEM data.

**Extended Data Fig. 8:**
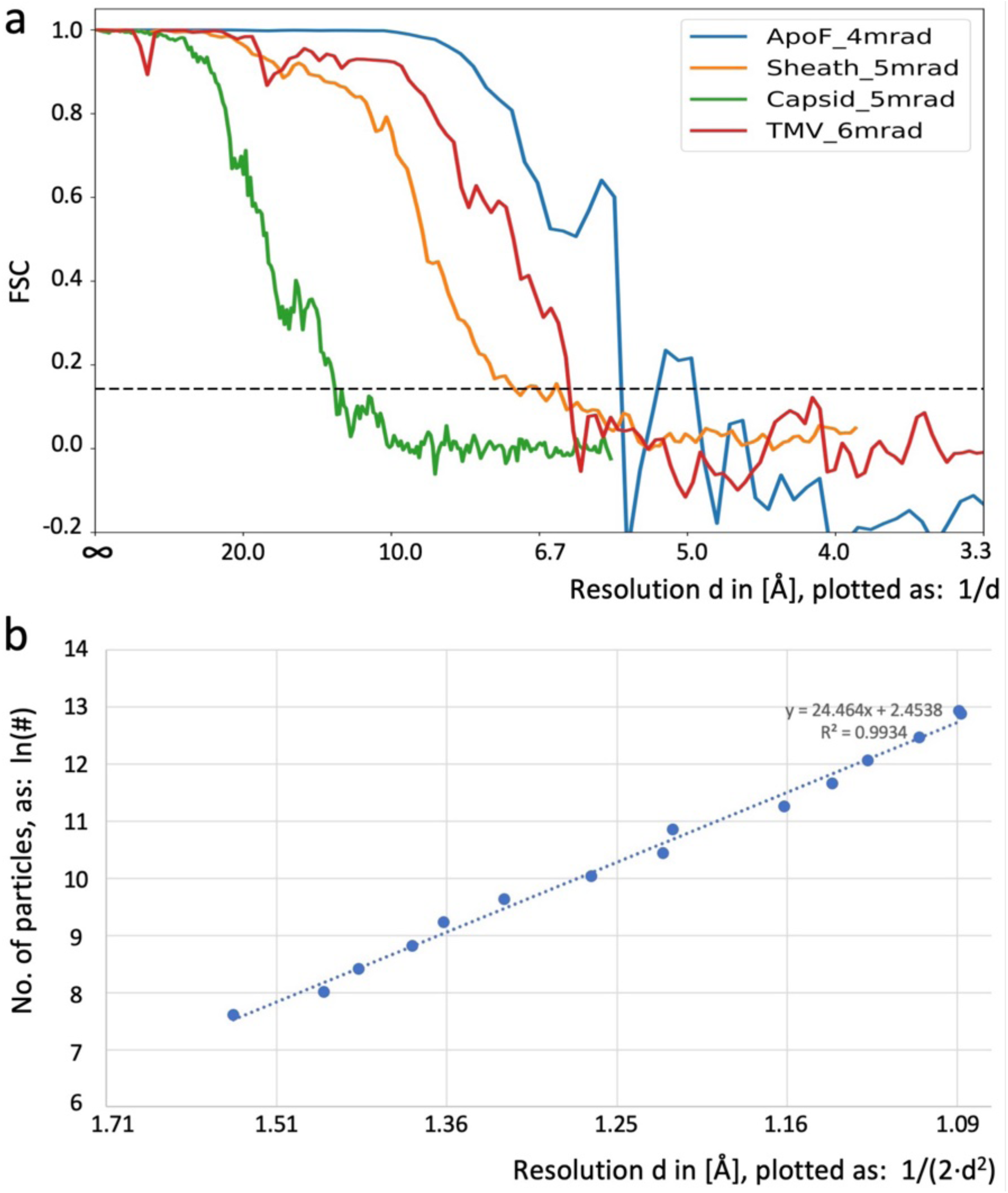
Resolution assessment of the determined structures. **a.** The Fourier Shell Correlation curves from the CryoSPARC processing of the 4D-STEM single particle datasets. **b.** The ResLog B-factor plot for the large ApoF cryo-EM dataset. Shown is the natural logarithm of the number of particles, plotted over the resolution (in 1/(2 * d^2^)). The slope of this plot indicates a ResLog value of 24.5 Å^-2^.

**Extended Data Fig. 9:**
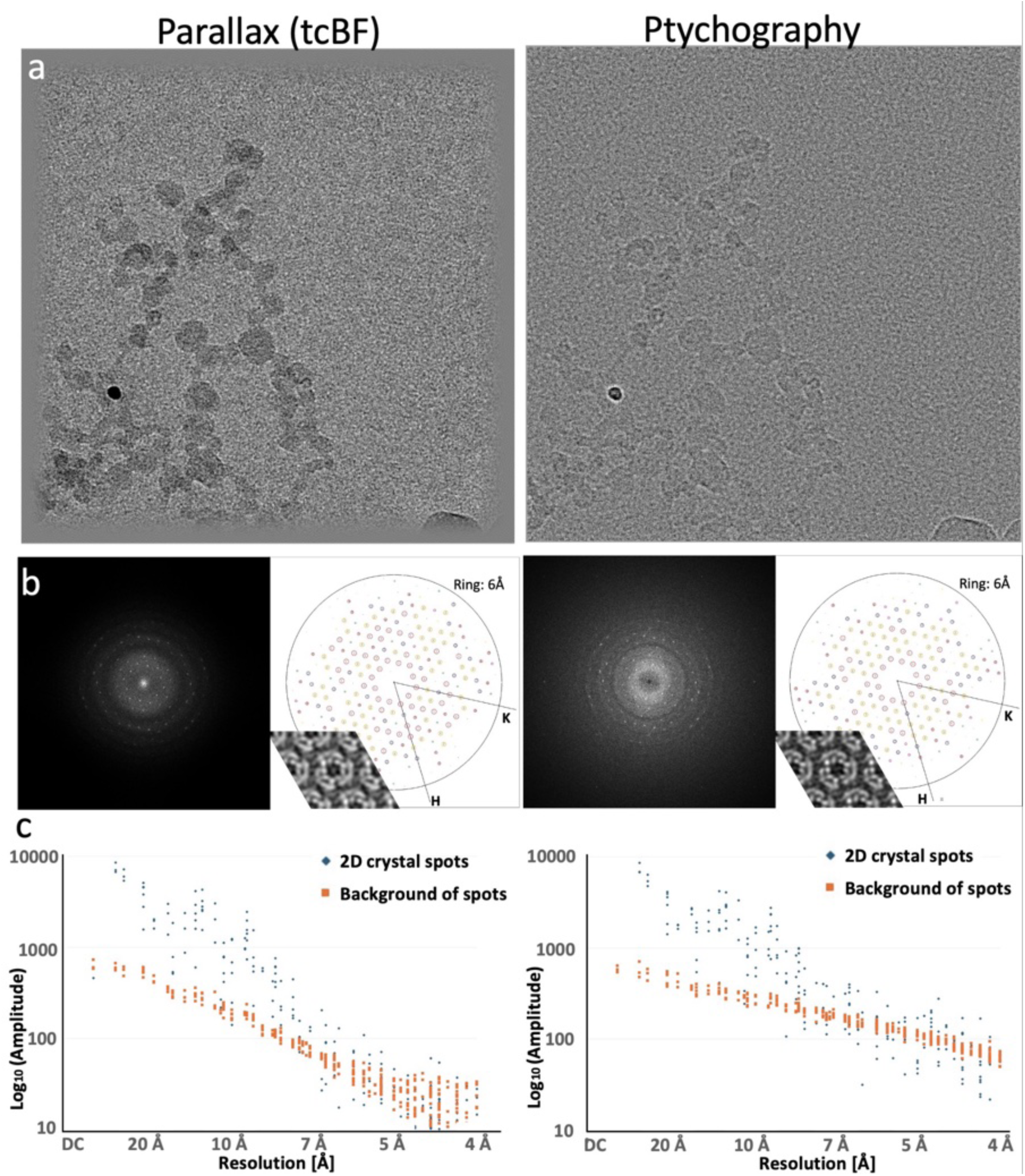
4D-STEM analysis of Bacteriorhodopsin 2D crystals. Glucose-embedded bacteriorhodopsin (BR) 2D crystals on continuous carbon film were imaged at LN2 temperature by 4D-STEM. **a.** Data processing with Parallax (tcBF, left) and with Ptychography (potential map, right) resulted in 2D images. **b.** Fourier transforms of the images shown in a (left), and IQ plot of diffraction spots (right). These indicate the SNR of each spot, calculated as the ratio of intensity-above-background divided by background. Background values are computed from the average of intensities on the edge of a 20×20 px square around each spot position. Inset: reconstructed projection map (2×2 unit cells). BR protein is white on a dark background. For the Parallax reconstruction, a Wiener filter CTF correction for a Scherzer CTF with 1.089 µm defocus, Cs=0mm, and 300kV was applied, which includes phase flipping. For the Ptychography reconstruction, no CTF correction was applied. **d**. Intensities above background and background of the 2D crystal spot, plotted on a logarithmic scale over the resolution of each spot.

**Extended Data Fig. 10:**
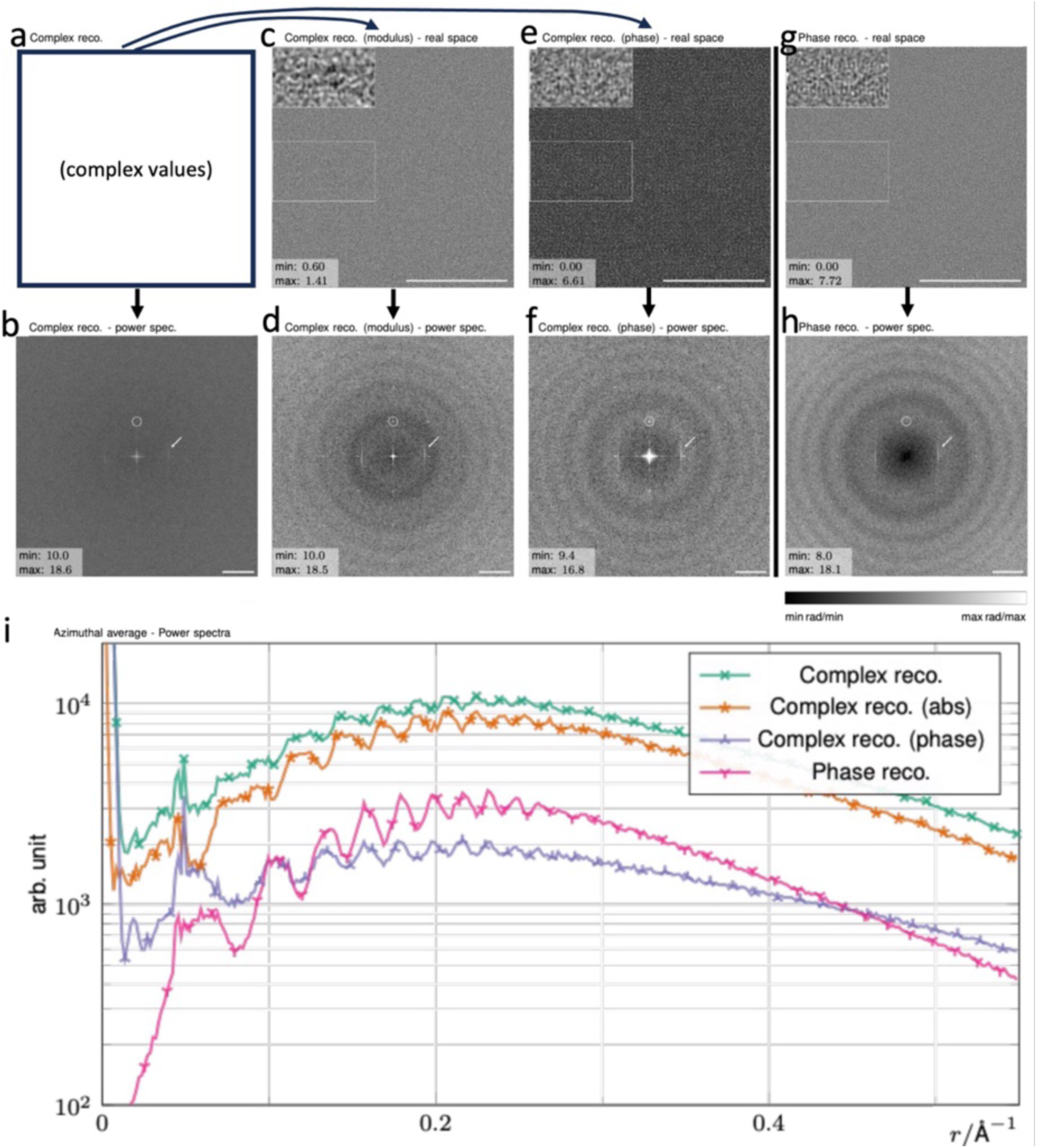
Impact of the object transmission function (OTF) model on the ptychography reconstruction. Using a 4D STEM dataset recorded at approx. 0.7 µm defocus of the TMV sample shown in Extended Data Fig. 7., two single-slice reconstructions were performed assuming a Poisson loss with a small regularization parameter in the logarithm to avoid divergence ^17^. **a** A complex object was reconstructed, so that both the modulus (i.e., amplitude, shown in **c**) and the phase (shown in **e**) of the OTF were reconstructed. **b.** shows the computed power spectrum of the complex reconstruction, which shows no ring-like pattern. **d.** shows the power spectrum of c, which shows a dark ring pattern. **e.** shows an image produced from the phase of the complex reconstruction. Its power spectrum is shown in **f. g.** shows a phase-only ptychography reconstruction. This was obtained by fixing the modulus (i.e., amplitude) of the complex reconstruction to constant “1”. Its power spectrum is shown in **h.** In the power spectra in b,d,f,h, the TMV layer lines are marked by an arrow, and artifacts from the scan pattern are marked with a white circle. The insets in c,e,g show Gaussian low-pass filtered versions of the white rectangles with standard deviations of 0.8. Scale bars in c,e,g = 100 nm. Scale bars in b,d,f,h = 0.43 nm^-1^. Note that the resulting phases of the full complex (**e,f**) and the pure phase (**g,h**) reconstruction differ only slightly, such that both are suitable as inputs for protein reconstructions. However, their power spectra exhibit rings in addition to the structural information. In particular, the Fourier transform of the full complex OTF in **b** does not show rings, as expected. Fourier transforming only the real-space modulus (**d**) or phase separately (**f**) introduces a ring pattern, which is also present in the Fourier transform of the pure phase reconstruction (**h)**. Note that the maxima in **d** match the minima in **f**. In **i,** radial averages of all power spectra are shown. These clearly reveal that the observed amplitude oscillations are not due to contrast inversions, which would be the case for the CTF of the classical defocused cryo-TEM.

## Supplementary Information

**Supplementary Table 1:**
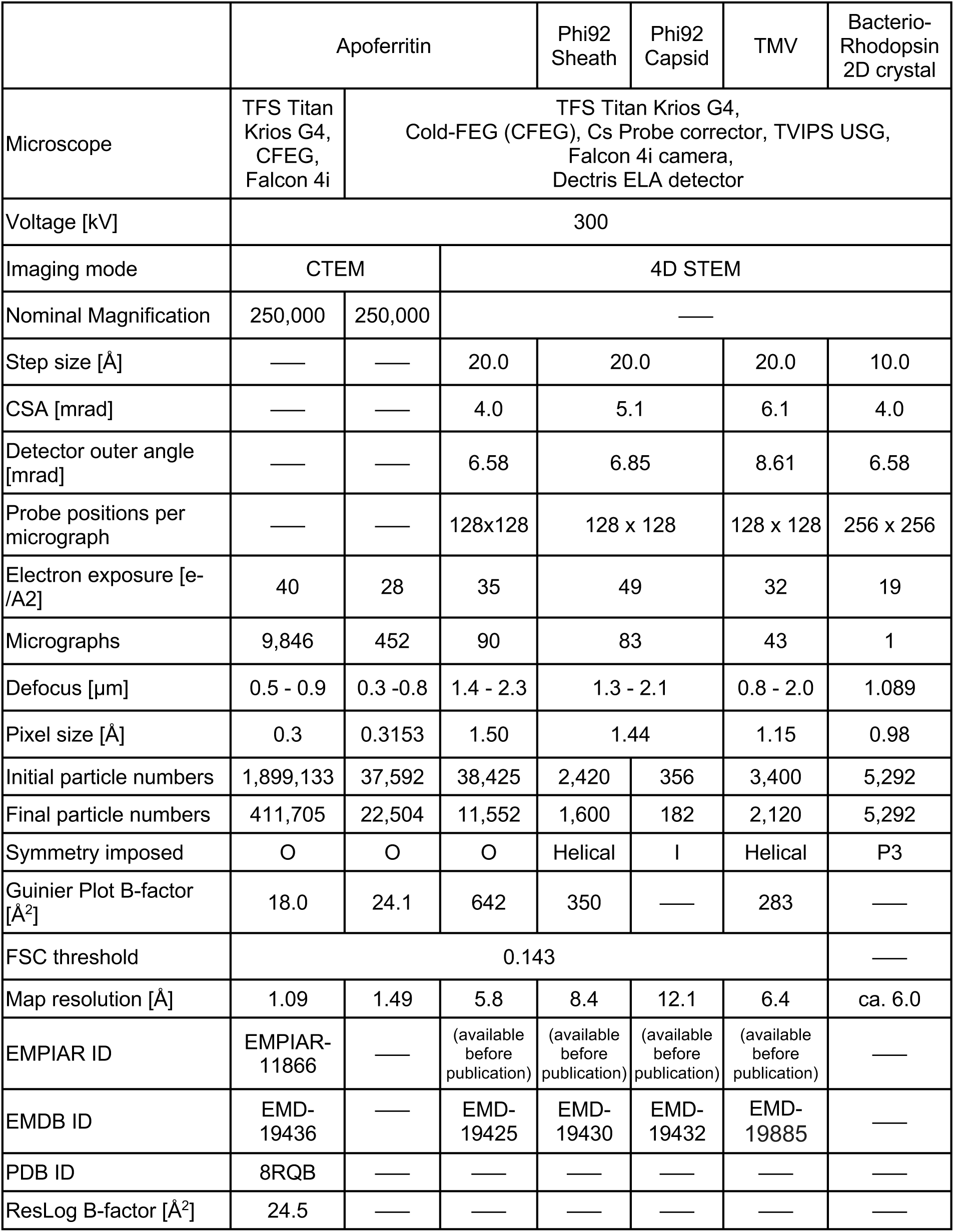
Cryo-EM and cryo-electron ptychography data collection and map statistics.

## References

1. Grant, T. & Grigorieff, N. Measuring the optimal exposure for single particle cryo-EM using a 2.6 A reconstruction of rotavirus VP6. Elife 4, e06980 (2015).

2. Brilot, A. F. et al. Beam-induced motion of vitrified specimen on holey carbon film. J. Struct. Biol. 177, 630–7 (2012).

3. Zhang, K. Gctf: Real-time CTF determination and correction. J. Struct. Biol. 193, 1–12 (2016).

4. Zheng, S. Q. et al. MotionCor2: anisotropic correction of beam-induced motion for improved cryo-electron microscopy. Nat. Methods 14, 331–332 (2017).

5. Danev, R. & Baumeister, W. Cryo-EM single particle analysis with the Volta phase plate. Elife 5, (2016).

6. Schwartz, O. et al. Laser phase plate for transmission electron microscopy. Nat. Methods 16, 1016–1020 (2019).

7. Lazić, I. et al. Single-particle cryo-EM structures from iDPC-STEM at near-atomic resolution. Nat. Methods 19, 1126–1136 (2022).

8. Evans, J. E. et al. Low-dose aberration corrected cryo-electron microscopy of organic specimens. Ultramicroscopy 108, 1636–1644 (2008).

9. Elad, N., Bellapadrona, G., Houben, L., Sagi, I. & Elbaum, M. Detection of isolated protein-bound metal ions by single-particle cryo-STEM. Proc. Natl. Acad. Sci. 114, 11139–11144 (2017).

10. Müller, S. A. & Engel, A. Mass measurement in the scanning transmission electron microscope: A powerful tool for studying membrane proteins. J. Struct. Biol. 121, 219–30 (1998).

11. Engel, A. Biological application of scanning probe microscopes. Annu. Rev. Biophys. Biophys. Chem. 20, 79–108 (1991).

12. Buban, J. P., Ramasse, Q., Gipson, B., Browning, N. D. & Stahlberg, H. High-resolution low-dose scanning transmission electron microscopy. J. Electron Microsc. (Tokyo) 59, 103–12 (2010).

13. Wolf, S. G. & Elbaum, M. CryoSTEM tomography in biology. Methods Cell Biol. 152, 197–215 (2019).

14. Rose, H. H. Future trends in aberration-corrected electron microscopy. Philos. Trans. Math. Phys. Eng. Sci. 367, 3809–3823 (2009).

15. Ophus, C. Four-dimensional scanning transmission electron microscopy (4D-STEM): From scanning nanodiffraction to ptychography and beyond. Microsc. Microanal. 25, 563–582 (2019).

16. Zhou, L. et al. Low-dose phase retrieval of biological specimens using cryo-electron ptychography. Nat Commun 11, 2773 (2020).

17. Diederichs, B., Herdegen, Z., Strauch, A., Filbir, F. & Müller-Caspary, K. Exact inversion of partially coherent dynamical electron scattering for picometric structure retrieval. Nat. Commun. 15, 101 (2024).

18. Chen, Z. et al. Electron ptychography achieves atomic-resolution limits set by lattice vibrations. Science 372, 826–831 (2021).

19. Rodenburg, J. M., McCallum, B. C. & Nellist, P. D. Experimental tests on double-resolution coherent imaging via STEM. Ultramicroscopy 48, 304–314 (1993).

20. Strauch, A. et al. Live Processing of Momentum-Resolved STEM Data for First Moment Imaging and Ptychography. Microsc. Microanal. 27, 1078–1092 (2021).

21. Rodenburg, J. M. & Bates, R. H. T. The theory of super-resolution electron microscopy via Wigner-distribution deconvolution. Philos. Trans. R. Soc. Lond. Ser. Phys. Eng. Sci. 339, 521– 553 (1997).

22. Bangun, A., Baumeister, P., Clausen, A., Weber, D. & Dunin-Borkowski, R. Wigner distribution deconvolution adaptation for live ptychography reconstruction. Microsc. Microanal. 29, 994– 1008 (2023).

23. Spoth, K. A., Nguyen, K. X., Muller, D. A. & Kourkoutis, L. F. Dose-Efficient Cryo-STEM Imaging of Whole Cells Using the Electron Microscope Pixel Array Detector. Microsc. Microanal. 23, 804– 805 (2017).

24. Varnavides, G., et al. Iterative Phase Retrieval Algorithms for Scanning Transmission Electron Microscopy. Preprint at 10.48550/arXiv.2309.05250 (2023).

25. Yu, Y., Spoth, K., Muller, D. & Kourkoutis, L. Dose-efficient tcBF-STEM imaging with real-space information beyond the scan sampling limit. Microsc. Microanal. 27, 758–760 (2021).

26. Yu, Y., Spoth, K., Muller, D. & Kourkoutis, L. Cryogenic TcBF-STEM Imaging of Vitrified Apoferritin with the Electron Microscope Pixel Array Detector. Microsc. Microanal. 26, 1736– 1738 (2020).

27. Elser, V. Phase retrieval by iterated projections. JOSA A 20, 40–55 (2003).

28. Maiden, A. M. & Rodenburg, J. M. An improved ptychographical phase retrieval algorithm for diffractive imaging. Ultramicroscopy 109, 1256–1262 (2009).

29. Hoppe, W. Elektronenbeugung mit dem Transmissions-Elektronenmikroskop als phasenbestimmendem Diffraktometer - von der Ortsfrequenzfilterung zur dreidimensionalen Strukturanalyse an Ribosomen. in *Angew*. Chem. vol. 95 465–494 (1983).

30. Yang, H., Pennycook, T. J. & Nellist, P. D. Efficient phase contrast imaging in STEM using a pixelated detector. Part II: optimisation of imaging conditions. Ultramicroscopy 151, 232–239 (2015).

31. Pei, X. et al. Cryogenic electron ptychographic single particle analysis with wide bandwidth information transfer. Nat. Commun. 14, 3027 (2023).

32. Yu, Y. et al. Dose-Efficient Cryo-Electron Microscopy for Thick Samples using Tilt-Corrected Scanning Transmission Electron Microscopy, Demonstrated on Cells and Single Particles. bioRxiv 2024.04.22.590491 (2024) doi:10.1101/2024.04.22.590491.

33. Mastronarde, D. N. SerialEM: A program for automated tilt series acquisition on Tecnai microscopes using prediction of specimen position. Microsc. Microanal. 9, 1182–1183 (2003).

34. Savitzky, B. H. et al. py4DSTEM: A Software Package for Four-Dimensional Scanning Transmission Electron Microscopy Data Analysis. Microsc. Microanal. 27, 712–743 (2021).

35. Yip, K. M., Fischer, N., Paknia, E., Chari, A. & Stark, H. Atomic-resolution protein structure determination by cryo-EM. Nature 587, 157–161 (2020).

36. Nakane, T. et al. Single-particle cryo-EM at atomic resolution. Nature 587, 152–156 (2020).

37. Punjani, A., Rubinstein, J. L., Fleet, D. J. & Brubaker, M. A. cryoSPARC: algorithms for rapid unsupervised cryo-EM structure determination. Nat. Methods 14, 290–296 (2017).

38. Scheres, S. H. RELION: implementation of a Bayesian approach to cryo-EM structure determination. J. Struct. Biol. 180, 519–30 (2012).

39. Zivanov, J., Nakane, T. & Scheres, S. H. W. Estimation of high-order aberrations and anisotropic magnification from cryo-EM data sets in RELION-3.1. IUCrJ 7, 253–267 (2020).

40. Henderson, R. & Unwin, P. N. Three-dimensional model of purple membrane obtained by electron microscopy. Nature 257, 28–32 (1975).

41. Biyani, N. et al. Focus: The interface between data collection and data processing in cryo-EM. J. Struct. Biol. 198, 124–133 (2017).

42. Leidl, M. L., Sachse, C. & Müller-Caspary, K. Dynamical scattering in ice-embedded proteins in conventional and scanning transmission electron microscopy. IUCrJ 10, 475–486 (2023).

43. O’Leary, C. M. et al. Contrast transfer and noise considerations in focused-probe electron ptychography. Ultramicroscopy 221, 113189 (2021).

44. Pelz, P. M. et al. Solving complex nanostructures with ptychographic atomic electron tomography. Nat. Commun. 14, 7906 (2023).

45. Hovden, R. et al. Breaking the Crowther limit: Combining depth-sectioning and tilt tomography for high-resolution, wide-field 3D reconstructions. Ultramicroscopy 140, 26–31 (2014).

46. Danev, R., Yanagisawa, H. & Kikkawa, M. Cryo-Electron Microscopy Methodology: Current Aspects and Future Directions. Trends Biochem. Sci. 44, 837–848 (2019).

47. Emsley, P., Lohkamp, B., Scott, W. G. & Cowtan, K. Features and development of Coot. Acta Crystallogr. D Biol. Crystallogr. 66, 486–501 (2010).

48. Yamashita, K., Palmer, C. M., Burnley, T. & Murshudov, G. N. Cryo-EM single-particle structure refinement and map calculation using Servalcat. Acta Crystallogr. Sect. Struct. Biol. 77, 1282– 1291 (2021).

49. Williams, C. J. et al. MolProbity: More and better reference data for improved all-atom structure validation. Protein Sci. 27, 293–315 (2018).

50. Rosenthal, P. B. & Henderson, R. Optimal determination of particle orientation, absolute hand, and contrast loss in single-particle electron cryomicroscopy. J. Mol. Biol. 333, 721–45 (2003).

